# Phenotypic pleiotropy of missense variants in human B cell-confinement receptor P2RY8

**DOI:** 10.1101/2025.02.28.640567

**Authors:** Taylor N. LaFlam, Christian B. Billesbølle, Tuan Dinh, Finn D. Wolfreys, Erick Lu, Tomas Matteson, Jinping An, Ying Xu, Arushi Singhal, Nadav Brandes, Vasilis Ntranos, Aashish Manglik, Jason G. Cyster, Chun Jimmie Ye

**Affiliations:** Division of Pediatric Rheumatology, Department of Pediatrics, University of California, San Francisco, San Francisco, CA, USA; Department of Microbiology and Immunology, University of California, San Francisco, San Francisco, CA, USA; Gladstone-UCSF Institute of Genomic Immunology, San Francisco, CA, USA; Department of Pharmaceutical Chemistry, University of California, San Francisco, San Francisco, CA, USA; Department of Epidemiology and Biostatistics, University of California, San Francisco, San Francisco, CA, USA; Howard Hughes Medical Institute, University of California, San Francisco, San Francisco, CA, USA; Division of Rheumatology, Department of Medicine, University of California, San Francisco, San Francisco, CA, USA; Institute for Human Genetics, University of California, San Francisco, San Francisco, CA, USA; Department of Biochemistry and Molecular Pharmacology, New York University, New York, NY, USA; Department of Bioengineering and Therapeutic Sciences, University of California, San Francisco, San Francisco, CA, USA; Bakar Computational Health Sciences Institute, University of California, San Francisco, San Francisco, CA, USA; Diabetes Center, University of California, San Francisco, CA, USA; Chan Zuckerberg Biohub, San Francisco, CA, USA; Quantitative Biosciences Institute, San Francisco, CA, USA; Department of Anesthesia and Perioperative Care, University of California, San Francisco, San Francisco, CA, USA; Parker Institute for Cancer Immunotherapy, University of California, San Francisco, San Francisco, CA, USA; Arc Institute, Palo Alto, CA, USA

## Abstract

Missense variants can have pleiotropic effects on protein function and predicting these effects can be difficult. We performed near-saturation deep mutational scanning of P2RY8, a G-protein-coupled receptor that promotes germinal center B cell confinement. We assayed the effect of each variant on surface expression, migration, and proliferation. We delineated variants that affected both expression and function, affected function independently of expression, and discrepantly affected migration and proliferation. We also used cryo-electron microscopy to determine the structure of activated, ligand-bound P2RY8, providing structural insights into the effects of variants on ligand binding and signal transmission. We applied the deep mutational scanning results to both improve computational variant effect predictions and to characterize the phenotype of germline variants and lymphoma-associated variants. Together, our results demonstrate the power of integrating deep mutational scanning, structure determination, and in silico prediction to advance the understanding of a receptor important in human health.

---

Missense variants, in which the amino acid sequence of a protein is altered, represent a crucial class of genetic variation that can significantly impact protein function. Germline missense variants are frequent in the human population^1,2^, highly enriched for those causal for rare Mendelian disorders, and statistically associated with common complex diseases^3^. Somatic missense mutations are frequent in cancers and those of driver genes are a major contributor to tumorigenesis^4,5^. Determining the effects of missense variants on protein function is essential for diagnosing and understanding the genetic causes of human diseases^2,6^.

Current approaches to missense variant annotation rely on both computational and experimental methods. Emerging computational variant effect prediction (VEP) approaches using deep learning can predict the effects of all missense mutations and have demonstrated success in identifying variants that cause Mendelian diseases, most of which are loss-of-function (LoF)^6–8^. However, these algorithms are much less effective at identifying cancer driver mutations, which are frequently gain-of-function (GoF) and not conserved evolutionarily^9,10^. Moreover, these tools provide only a single functional prediction per variant, limiting their capacity to predict effects across diverse phenotypes.

In parallel, the convergence of next-generation sequencing and massively-parallel gene synthesis have propelled experimental profiling of large numbers of missense variants via deep mutational scanning (DMS), also known as multiplexed assays of variant effect (MAVE)^11–13^. Unlike computational predictions, DMS experiments can assess variant effects across multiple phenotypes and cell types, offering rich, direct data on functional consequences. Furthermore, because DMS does not depend on evolutionary conservation, they provide an unbiased approach to variant characterization. DMS approaches remain low-throughput, and most human genes have yet to be profiled. Nevertheless, the growing body of DMS data, including an international collaborative effort to collate these results^14^, represents a valuable resource that could ultimately inform and refine computational VEP tools.

One application of these approaches is studying G-protein-coupled receptors (GPCRs), a large family of ∼800 transmembrane receptors that play important roles across numerous physiologic pathways and are the most frequent drug targets in medicine^15,16^. Sensitivity to missense variants generally varies between domains within a GPCR, as illustrated by the uneven distribution of GPCR variants that cause Mendelian disorders^17^. Transducers of GPCR signaling include the heterotrimeric G proteins, of which there are four major families, G_s_, G_i/o_, G_q/11_, and G_12/13_, as well as the β-arrestins^18,19^. Several GPCRs have been the focus of DMS, revealing insights into how missense variants affect their surface expression^20–22^ and signaling^23,24^. Examining both expression and function has the benefit of allowing one to distinguish between effects mediated through, or independent of, changes in expression^24^. Despite this progress, GPCRs that signal primarily through G_12/13_ remain underexplored among DMS and structural studies^25^.

P2RY8 is a G_13_-coupled GPCR that is expressed on several lymphocyte subsets, including germinal center (GC) B cells, where it restrains cell migration and proliferation^26–28^. Its ligand is S-geranylgeranyl-L-glutathione (GGG)^29^. Addition of GGG in migration assays of P2RY8-expressing cells causes activation of RhoA and inhibition of migration towards chemokines^30^. The *P2RY8* locus is located on the pseudoautosomal region of the X and Y chromosomes and is frequently mutated in diffuse large B cell lymphoma (DLBCL) and Burkitt lymphoma^26,31,32^. In addition, variants in P2RY8 have been identified in a small number of patients with lupus, and these variants were found to affect B cell negative selection and plasma cell development when expressed in mice^33^.

The immunologic importance of P2RY8 and its restricted expression make it an attractive target for therapeutic intervention, but the current understanding of how specific missense variants affect its function is limited. To address this, we conducted DMS of P2RY8, evaluating surface expression and two distinct functional outcomes, inhibition of migration and restraint of proliferation. Using cryo-electron microscopy (cryo-EM), we determined the structure of P2RY8 in complex with its ligand, GGG, providing a structural scaffold for interpreting how missense variants influence its active conformation and signaling. We also illustrate how integrating relatively sparse experimental data from these screens can enhance the performance of computational VEP tools. We performed in-depth validation of select variants, which provided further insights into P2RY8 function, including evidence that receptor initiation of pathways regulating migration and proliferation is at least partly distinct. In sum, we demonstrate the utility of performing DMS on multiple functional phenotypes and the complementary benefits of a multimodal approach to annotate the pleiotropic effects of missense variants of a GPCR to better understand its biology.

## Results

### Deep mutational scanning of P2RY8 across three phenotypes

We performed deep mutational scanning of P2RY8, using a pooled lentiviral library of 7,045 variants covering all possible substitutions for 353/358 non-start codons and synonymous variants for 338 positions (Supplemental Table 1). We transduced OCI-Ly8 (Ly8) cells, a human DLBCL cell line, in which we previously knocked out endogenous P2RY8^29^, and performed three parallel screens: (1) receptor surface expression, (2) migration of cells simultaneously exposed to chemoattractant CXCL12 and P2RY8 ligand GGG, and (3) proliferation across multiple time points over 10 days (Fig. 1a and Extended Data Fig. 1a,b). Each of these three screens were highly reproducible across four replicates, with pairwise Pearson correlation coefficients (r) ranging from 0.96 to 0.97 for surface expression, 0.71 to 0.85 for migration, and 0.33 to 0.66 for proliferation (Extended Data Fig. 1c-e). Missense variants showed a bimodal distribution of effect sizes (i.e., z-scores based on synonymous variant effects) across all three assays (Fig. 1b,c and Extended Data Fig. 2a,b and Supplemental Table 2). Approximately half of missense variants reduced the expression of surface P2RY8 but 365 variants increased expression (Fig. 1c), suggesting GoF effects are detectable in our screens. As expected, the effect sizes of variants on expression of P2RY8 are inversely correlated with their effects on migration (Fig. 1c; Spearman correlation coefficient (ρ) of −0.837) and proliferation (Extended Data Fig. 2a, ρ = −0.614). The migration and proliferation effect sizes were also significantly correlated (Extended Data Fig. 2b, ρ = 0.725).

**Fig. 1:**
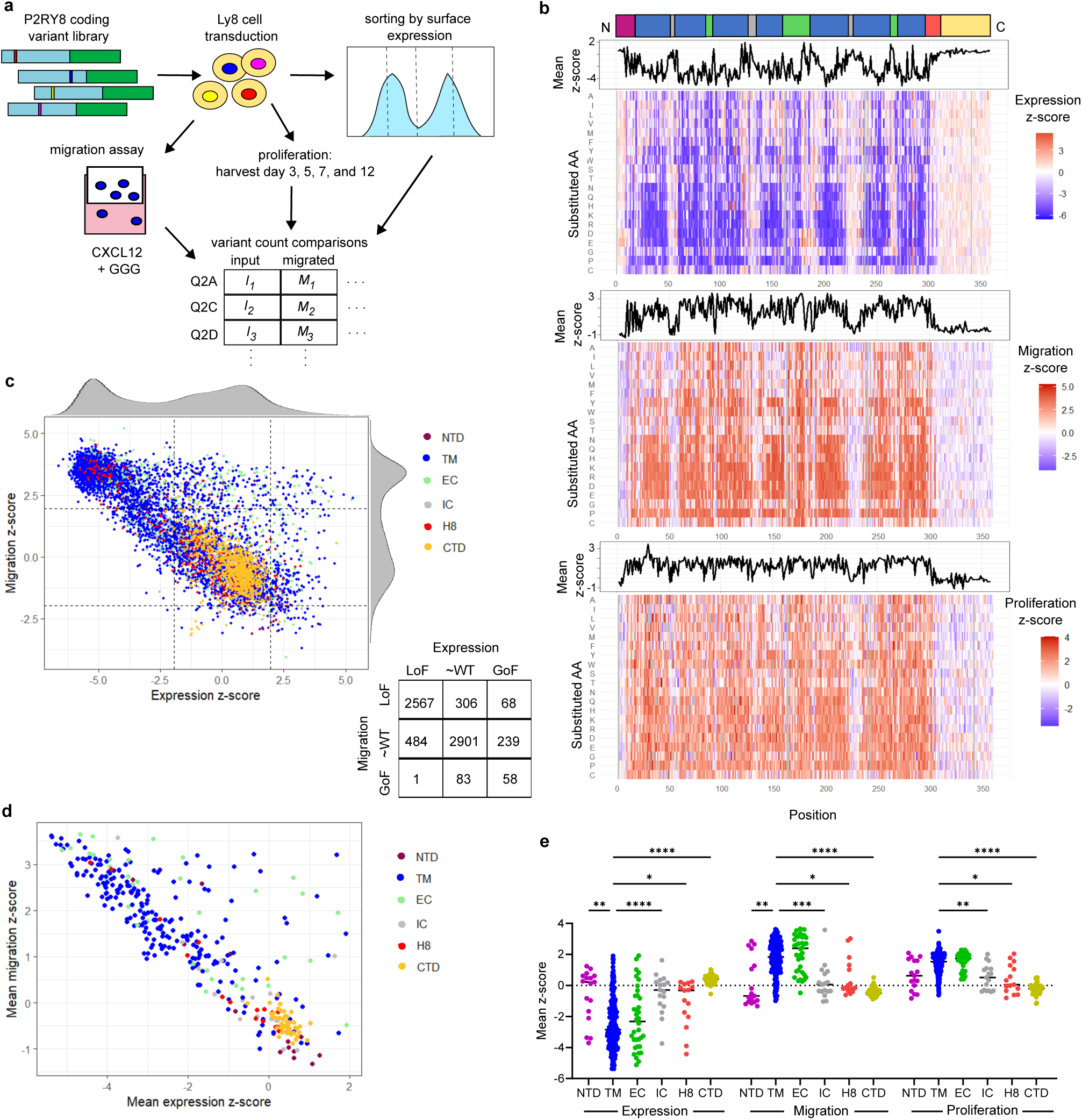
DMS of P2RY8 across three phenotypes. **a,** Schematic of DMS approach **b,** Heat maps of surface expression, migration, and proliferation z-scores with line plots of mean z-score for each position, **c,** Plot comparing expression and migration z-scores of missense variants, colored by protein domain, with table showing variant count by section, boundaries at z-scores of 2 and - 2. **d,** Plot comparing expression and migration mean z-scores for each position, colored by protein domain. **e,** Expression, migration, and proliferation mean z-scores for each position, partitioned by protein domain; lines are medians. Negative z-scores are LoF for expression, GoF for migration or proliferation. Welch ANOVA test with Dunnett’s T3 multiple comparisons test; for each assay TM compared to the five other domains; adjusted p-value * < 0.05, ** <0.01, *** < 0.001, **** < 0.0001. C, C-terminus; CTD, C-terminal domain; EC, extracellular loop; GoF, gain-of-function; H8, helix 8; IC, intracellular loop; LoF, loss-of-function; N, N-terminus; NTD, N-terminal domain; TM, transmembrane domain.

Averaging the effect sizes across substitutions for each position resulted in higher inverse correlation between migration and expression while also identifying several positions in which missense variants consistently affected function but not expression (Fig. 1d and Extended Data Fig. 2c,d). Mutation tolerance at specific positions varied significantly within each protein domain; highly sensitive positions were enriched in the transmembrane helices (TM) and extracellular loops (EC), whereas no position in the C-terminal domain (CTD) was highly sensitive (Fig. 1e). Similar differences between domains were also observed when examining the effect of individual variants, though large effect substitutions do exist in all domains (Extended Data Fig. 2e). The range of variant effects within positions varied widely, with some positions having similar effects for all missense variants and others having a large spread of effects (Extended Data Fig. 2f,g).

### Improving computational variant effect prediction using limited experimental data

ESM1b is a self-supervised protein language model that has been shown recently to have high zero-shot predictive power of pathogenicity of coding variants^6^. We found that our DMS expression effect sizes correlated well overall with the variant ESM1b scores (Fig 2a, ρ = 0.586). However, there was a subset of variants in which decreased expression was not successfully predicted (Fig. 2a). There was similar correlation between ESM1b scores and and migration effect sizes but lower correlation with proliferation effect sizes (Fig. 2b,c; ρ = −0.593 for migration and ρ = −0.500 for proliferation). When averaged by position the mean effect sizes from the DMS were better correlated with mean ESM1b scores for all three assays (Fig. 2d-f). Correlation between ESM1b and the DMS results varied by protein domain, with consistently poor correlation within the CTD (Fig. 2g). Consistent with previous reports, we found that ESM1b pathogenicity scores did not reliably distinguish between WT-like and GoF variants (Fig. 2h).

**Fig. 2:**
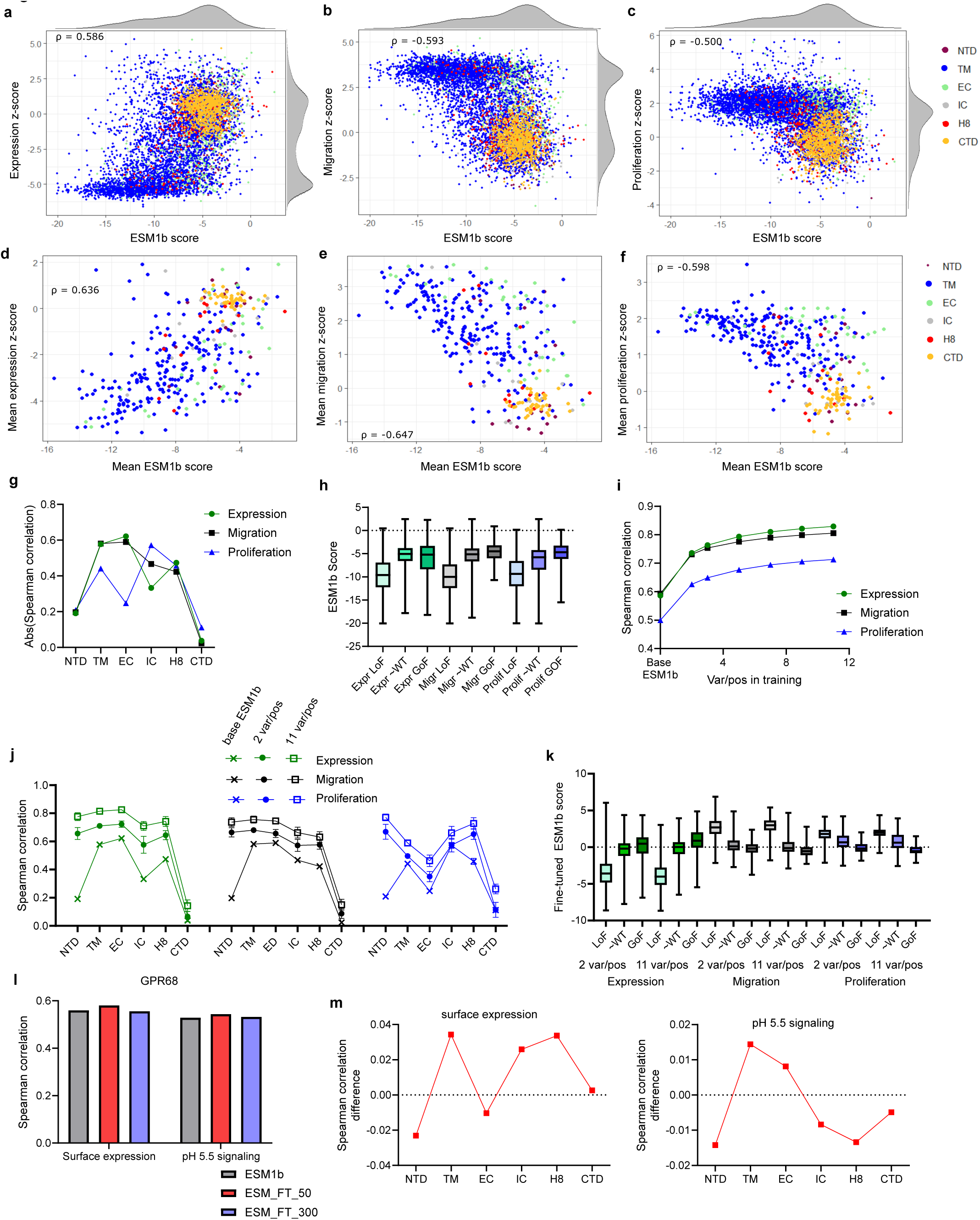
Improving computational variant effect prediction using limited experimental data. **a-c** Plots comparing ESM1b scores and **(a)** expression z-score, **(b)** migration z-score, or **(c)** proliferation z-score for each variant, colored by protein domain. **d-f** Plots comparing mean ESM1b score and **(d)** expression mean z-score, **(e)** migration mean z-score, or **(f)** proliferation mean z-score for each position, colored by protein domain. **g,** Plot of Spearman correlation between ESM1b score and variant expression, migration, and proliferation z-scores, partitioned by protein domain. **h,** Plot comparing distribution of ESM1b scores for each variant, partitioned by DMS phenotype. Whiskers extend to maximum and minimum, boxes shows 25th percentile, median, and 75th percentile. **i,** Plot of Spearman correlation between fine-tuned ESM1b scores and DMS z-scores as training set size varies. Shows mean and SD (k = 50). **j,** Plot comparing Spearman correlation when using ESM1b score or fine-tuned ESM1b scores with 2 variants/position or 11 variants/position training sets, partitioned by domain. Shows means and SD (k = 50). **k,** Plot comparing distribution of fine-tuned ESM1b scores for representative 2 variant per position training and representative 11 variant per position training, partitioned by DMS phenotype. Whiskers extend to maximum and minimum, boxes shows 25th percentile, median, and 75th percentile. **l,** Plot comparing Spearman correlations between human GPR68 DMS results^24^ and base ESM1b or ESM1b fine-tuned on P2RY8 expression results for 50 or 300 fine-tuning steps. **m,** Plots showing difference in Spearman correlation between base ESM1b and 50-step fine-tuned ESM1b on GPR68 surface expression (L) or pH 5.5 signaling (R) (value > 0 indicates greater correlation with fine-tuning). In **g, i, j,** sign inverted for migration and proliferation so all three correlations have same sign. CTD, C-terminal domain; EC, extracellular loop; GoF, gain-of-function, z-score > 2 for expression, < −2 for migration, proliferation; H8, helix 8; IC, intracellular loop; LoF, loss-of-function, z-score < −2 for expression, > 2 for migration, proliferation; NTD, N-terminal domain; TM, transmembrane helices; ρ, Spearman correlation coefficient.

We investigated whether fine-tuning ESM1b with limited amounts of training data from our DMS screen would yield a better supervised predictor of DMS results. Across training sets of 2, 3, 5, 7, 9, and 11 variants per position, we observed a sharply increased correlation between predicted scores and the DMS effect sizes with just 2 variants per position, with further improvements with increasing variants per position (Fig. 2i). For example, for the expression data, correlation was ρ = 0.586 with base ESM1b scores but the mean ρ was 0.736 after training with 2 variants per position (∼10% of the DMS data), 0.794 with 5 variants per position, and 0.829 with 11 variants per position. Across all three phenotypes, 2 variants per position produced ∼60% as much improvement as was achieved with 11 variants per position. The fine-tuning included a final ridge regression step as we observed that this resulted in greater increases in correlation, particularly with smaller training sets (Extended Data Fig. 3a).

Given the significant overall gains observed even with 2 variants per position, we then examined the degree to which domain-specific changes were observed. We observed the most striking improvement within the NTD and the smallest amount of improvement within the CTD, with intermediate improvements for other domains (Fig. 2j). Training with 11 variants per position showed further improvements though correlation in the CTD remained modest (Fig. 2j). The influence of the base ESM1b remained evident, with domains with higher correlations with base ESM1b continuing to be the domains with higher correlations after fine-tuning, with the exception of the NTD, for which base ESM1b was poorly predictive but even modest amounts of training data resulted in a large improvement.

The improved correlation was visible when plotting fine-tuned ESM1b and DMS expression effect sizes relative to plotting with baseline ESM1b (Extended Data Fig. 3b, relative to Fig. 2a). Compared with what was seen with base ESM1b, there was marginally better separation between the WT-like and GoF variants with 2 variants per position training, with improved separation with 11 variants per position, though some overlap remained (Fig. 2k). These findings suggest that although a screen with as few as two variants per position can enhance predictions of variant effects, identification of GoF variants will likely require more intensive experimental analysis, barring the development of new, superior VEP algorithms.

To ensure these observations were not unique to ESM1b, we also compared our DMS results with an alternative VEP tool, AlphaMissense (AM). AM was produced through fine tuning AlphaFold on human and primate population variant frequencies and has been shown to have state-of-the-art zero-shot performance at predicting DMS results^7,34^. We observed high correlation between AM scores and the expression, migration, and proliferation effect sizes, in fact with higher correlation in each case than for ESM1b (Extended Data Fig. 4a-c). Similarly, correlation was again higher for the position-averaged data compared to individual variant data (Extended Data Fig. 4d-f). As with ESM1b, correlation between AM and the DMS data varied by protein domain, with much lower correlation with the CTD (Extended Data Fig. 4g). As with ESM1b, AM better distinguished between LoF variants and WT-like variants than between GoF variants and WT-like variants (Extended Data Fig. 4h). LoF variants that had a benign AM pathogenicity score^7^ were overrepresented in the EC, IC, and CTD domains relative to correctly assigned LoF variants (Extended Data Fig. 4i). In addition, these misclassified variants were overrepresented at positions in which the WT amino acid is positively charged and among variants in which the substituted amino acid was glycine or an aliphatic hydrophobic residue (Extended Data Fig. 4i).

Fine-tuning comparable to what was performed with ESM1b was not possible with AM given the specific model weights are not publicly available. However, concurrently with the ESM1b fine-tuning described above, we also implemented the augmented one-hot encoded (OHE) ridge regression model previously described^35^, using the same variant training sets as in the fine-tuning iterations. This resulted in improved correlation between predicted scores and the DMS effect sizes, with greater gains with increasing training set size (Extended Data Fig. 4k). The degree of improvement was less than was seen with ESM1b fine-tuning, though given the higher baseline correlation, the absolute correlations differences were smaller between the two (Extended Data Fig. 3c). The improvements with OHE-regression with ESM1b were also less than those seen from fine-tuning ESM1b, particularly with larger training sets (Extended Data Fig. 3d). In summary, two different approaches using a subset of our DMS data were able to improve variant effect prediction relative to two distinct state-of-the-art VEP tools alone.

Finally, we investigated whether the DMS results for P2RY8 could be used to improve variant effect prediction for another GPCR. Recently a DMS study was published for GPR68, a pH sensing receptor that like P2RY8 is a class A, subgroup δ GPCR^24^. The correlations between ESM1b scores and GPR68 DMS scores for missense variants were ρ = 0.560 for surface expression and ρ = 0.529 for pH 5.5 signaling, comparable to what was seen with P2RY8. We found that after fine-tuning ESM1b on P2RY8 DMS expression with 300 optimization steps, there was no improvement in variant effect prediction for GPR68 (Fig. 2l), with a Spearman correlation difference of <0.005 for both GPR68 expression and function. We restricted to 50 optimization steps to reduce overfitting and now observed a small increase in correlation of ∼0.02, in comparison to ∼0.13-0.15 increase within P2RY8 itself after training with 2 variants and ∼0.21-0.24 increase after training with 11 variants (Fig. 2j). We did observe varying effects by protein domain, with the greatest improvement in the TM domain for expression and function, with significant gains in ICL and H8 regions for expression as well (Fig. 2m). This finding corresponds to greater conservation of these regions across GPCRs^36,37^.

In summary, use of a subset of our DMS data was able to strongly improve variant effect prediction relative to two distinct state-of-the-art VEP tools alone, but showed limited ability to generalize to improvements in variant effect predictions in a different GPCR.

### Validation of variants with independent effects on expression and function

Although there was overall inverse correlation between variant effects on cell migration and surface expression, we identified several hundred variants that affected migration independent of expression but not vice versa, consistent with the hierarchical nature of these phenotypes (Fig. 1c). We computed an expression-adjusted migration score, and identified 25 residues with substantially higher mean effect size across substitutions in migration than in expression; 23/25 of these positions were located within a transmembrane helix or extracellular loop (Fig. 3a). Plotting the expression-adjusted migration score by position across the protein sequence showed several peaks, including in TM3 into ICL2, in ECL2, and in TM6 into ECL3 (Fig. 3b). The first region includes the canonical D(E)-R-Y motif that is critical for GPCR signaling^18^. These 25 positions also generally had larger mean effect sizes on proliferation than expression (Extended Data Fig. 5a). Performing a comparable analysis on migration and proliferation, there were only six positions with a high migration-adjusted proliferation scores and none with the converse (Fig. 3c).

**Fig. 3:**
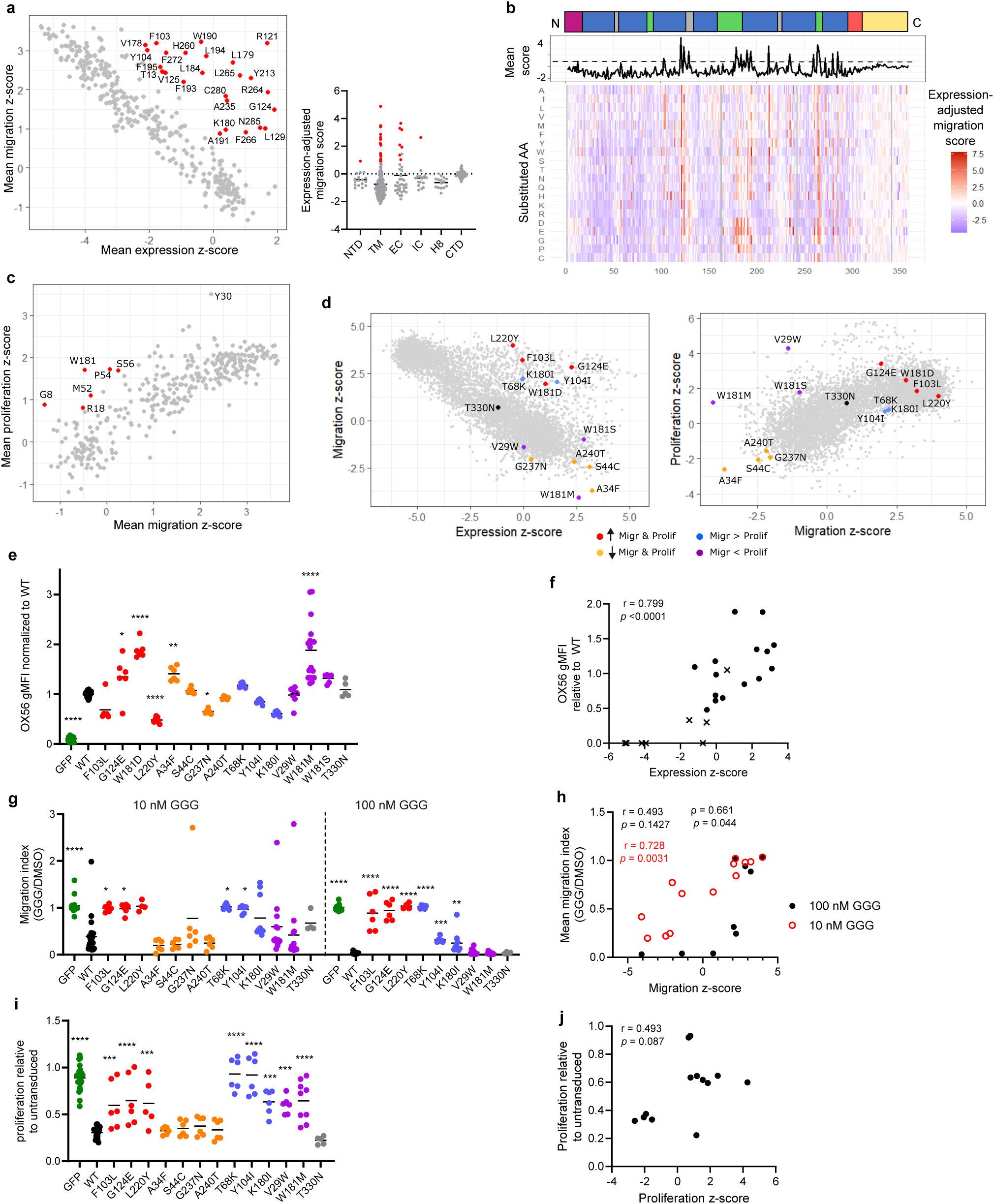
Validation of variants with independent effects on expression and function. **a,** (L) Plot comparing expression and migration mean z-scores for each position, those with high expression-adjusted migration scores are colored red; (R) Expression-adjusted migration score for each position, partitioned by domain, lines are medians. **b,** Heat map of expression-adjusted migration score for each variant, as well as line plot of mean expression-adjusted migration scores by position. **c,** Plot comparing migration and proliferation mean z-scores for each position, those with high migration-adjusted proliferation score are colored red. **d,** Plots comparing expression and migration z-scores (L) or migration and proliferation z-scores (R), with select variants labeled. **e,** P2RY8 surface expression normalized to WT P2RY8, lines are means. Pooled results from 9 experiments, total 5 to 26 biological replicates per condition. **f,** Plot of DMS expression z-score and WT-normalized expression for select variants, circles are variants in **d**, crosses are variants assayed previously^26,29^. **g,** Migration of cells toward CXCL12 in the presence of GGG, relative to DMSO (vehicle); each dot is a biological replicate, lines are means. Pooled results from 9 experiments, total 4 to 23 biological replicates per condition. **h,** Plot of DMS migration z-score and migration indices for variants shown in **g. i,** Proliferation over 13 days relative to untransduced cells in same well; each point is a biological replicate, lines are means. Pooled results from 8 experiments, total 5 to 21 biological replicates per condition. **j,** Plot of DMS proliferation z-score and proliferation relative to untransduced cells for variants shown in **i**. **e, g, i,** One-way ANOVA test with Dunnett’s multiple comparisons test, each variant compared to WT; adjusted p-value * < 0.05, ** <0.01, *** <0.001, **** < 0.0001. **f, h, j,** Two-tailed p-values; r, Pearson correleation coefficient; ρ, Spearman correlation coefficient.

We performed validation studies of several variants, drawing from different regions of the protein and representative of various migration and proliferation phenotypes with preserved surface expression: increased migration and proliferation (F103^3×32^L, G124^3×53^E, W181^ECL2^D, L220^5×65^Y), decreased migration and proliferation (A34^1×43^F, S44^1×53^C, G237^6×35^N, A240^6×38^T), increased migration relative to proliferation (T68^2×49^K, Y104^3×33^I, K180^ECL2^I), increased proliferation relative to migration (V29^1×38^W, W181^ECL2^M, W181^ECL2^S), and WT-like function (T330^CTD^N) (Fig. 3d). (Superscripts refer to generic GPCR residue numbering per revised Ballesteros-Weinstein method for class A GPCRs^36,38^.) These variants include four of the positions with high expression-adjusted migration scores (Fig. 3a) and one of the residues with high migration-adjusted proliferation scores (Fig. 3c). All variants were confirmed to support surface expression, and high correlation was observed between screen and validation results for these 15 variants (Fig. 3e,f; r = 0.506 with p-value = 0.0543). The inclusion of 8 additional P2RY8 variants that we have previously characterized^26,29^ resulted in a higher correlation (r = 0.799, p-value < 0.0001). Despite P2RY8 expression, cells with these variants showed a range of migratory capacity toward CXCL12 in the presence of GGG (Fig. 3g). We observed high correlation between the screen and validation at both high and low GGG exposure (100 nM, r = 0.493 with p-value = 0.1427, though Spearman ρ = 0.661 with p-value = 0.044; 10 nM, r = 0.728, p-value = 0.0031) (Fig. 3h). We also performed validation proliferation studies, comparing change in relative abundance of transduced and untransduced cells over a 13-day span. Over that time, WT P2RY8 results in ∼2/3 reduction in frequency compared to untransduced cells. Although the effects of the variants on proliferation appeared largely in agreement with those seen in the screen, the trend was not statistically significant (Fig. 3i,j). In summary, individual variant validation across all three assays yielded findings that were in close accord with the screen.

### Structure of activated ligand-bound human P2RY8

To facilitate understanding of the structural basis of the variant phenotypes revealed by the DMS, we used cryo-EM to determine the structure of activated, GGG-bound human P2RY8. We generated a C-terminal fusion of P2RY8 with a minimized and stabilized version of Gα_13_ (miniG_13_) as previously described for other heterotrimeric G proteins^39,40^ (Extended Data Fig. 6a). P2RY8-miniG_13_ was purified in the presence of GGG and further complexed with recombinant Gβ1γ2 (Extended Data Fig. 6b). The resulting preparation was analyzed by single-particle cryo-EM, which yielded a 2.7 Å map of the complex (Extended Data Fig. 7 and Supplemental Table 3). While the receptor and miniG_13_ are well resolved, only a portion of the Gβ_1_ was visible in the resulting density; the majority of Gγ_2_ was not resolved. Importantly, the map revealed a density consistent with GGG located in the canonical class A GPCR orthosteric ligand-binding pocket (Fig. 4a,b).

**Fig. 4:**
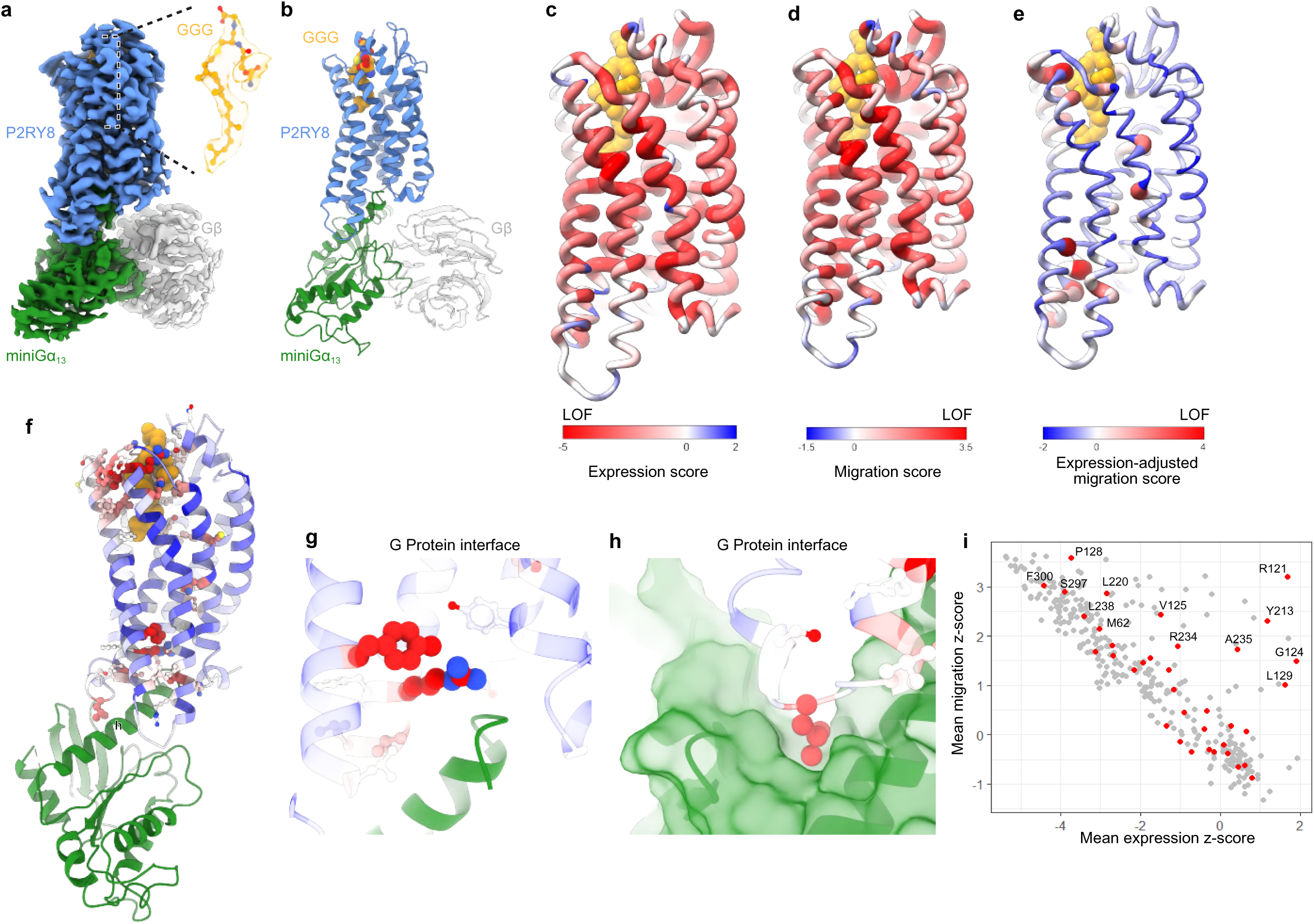
Structure of activated ligand-bound human P2RY8. **a,b,** Cryo-EM density map **(a)** and ribbon model **(b)** of active human P2RY8 bound to GGG (orange). P2RY8 is fused to mini-Gα_13_ (green) and bound to G*β*_1_ (grey). **c, d,e,** Ribbon model of P2RY8 colored to indicate DMS mean effect sizes for expression **(c)**, migration **(d)**, and expression-adjusted migration score **(e)**. Grey indicates positions with no data available. **f, g, h,** Expression-adjusted migration scores for individual residues with close-up views of **(g)** D(E)-R-Y motif and **(h)** G-protein interface. **i,** Plot comparing expression and migration mean z-scores by position, with Gα_13_-interacting positions colored red.

We next mapped DMS functional results onto the receptor structure, which illustrated the structural elements most essential for expression and migration inhibition (Fig. 4c,d). As observed in scatter plots of the DMS data (Fig. 1d), there was substantial correlation between P2RY8 residues required for expression and migration inhibition. Variants of positions that lead to decreased P2RY8 expression are primarily concentrated in the TM regions, with EC and IC loops more tolerant to variants. Similarly, most TM variants cause decreased migration inhibition. To identify positions important for migration that do not simply affect protein expression, we also mapped expression-adjusted migration scores (Fig. 4e). This analysis revealed two critically important regions: residues surrounding the GGG ligand and key interacting residues with Gα_13_ (Fig. 4f).

The P2RY8 interaction with miniGα_13_ is highly similar to the canonical conformation observed for most GPCR-G protein complexes to date^37,41^. Our DMS provides an unbiased view of key regions of P2RY8 that are important for signal transduction. We observed that six residues directly contacting miniGα_13_ had high expression-adjusted migration scores: R121^3×50^, G124^3×53^, V125^3×54^, L129^34×51^, Y213^5×58^, and A235^6×33^ (Fig. 3a and 4f), and two reisdues had high migration-adjusted proliferation scores: M52^ICL1^ and S56^2×37^. Although we lack an inactive state structure of P2RY8, the importance of these residues is underscored by their conservation among class A GPCRs and key contacts they make to stabilize P2RY8 or interact with Gα_13_. For example, both R121^3×50^ and Y213^5×58^ directly bind to the α5 helix of miniGα_13_ (Fig. 4g). As has been observed previously for many class A GPCRs, Y213^5×58^ engages Y293^7×53^ in a bonding network that likely stabilizes active P2RY8. Additionally, L129^34×51^ binds to a conserved hydrophobic cavity in Gα_13_ (Fig. 4h). Plotting the average DMS effect sizes for all 38 P2RY8 positions in contact with miniGα_13_ showed a range of mutation tolerance ranging from highly sensitive for both expression and migration to largely tolerant (Fig. 4i). Individual missense variants for two of these positions, G124^3×53^E and L220^5×65^Y, were including in the variant validation set already described; both of these variants were expressed but strongly deleterious to function (Fig. 3e,g,i). Taken together, the DMS results and cryo-EM structure provide complementary, concordant insights into receptor function.

### Structural features of GGG recognition

GGG binds to P2RY8 in a large solvent-facing cavity, with a combination of hydrophilic interactions with the glutathione headgroup and hydrophobic interactions along the geranylgeranyl tail (Fig. 5a and Extended Data Fig. 8). The glutathione headgroup is located towards the extracellular side of the pocket, while the geranylgeranyl tail extends deeply within the core of the receptor. Of the 28 residues that contact GGG, ten of the residues are among those with high expression-adjusted migration scores (Fig. 3a and 5b). Several of these high scoring positions contact the GGG headgroup, including K180^ECL2^, H260^6×58^, R264^ECL3^, and Y272^7×32^. Residues H260^6×58^, R264^ECL3^, and Y272^7×32^ engage the glutamate residue in GGG, with H260^6×58^ making a direct hydrogen bond with the carboxylic acid. Additionally, several positions contact the proximal geranylgeranyl tail closest to the thioether cysteine in GGG; these include L179^ECL2^, W190^5×35^, L194^5×39^, and L265^ECL3^. In addition, two residues in contact with the GGG headgroup, G8^NTD^ and W181^ECL2^, have high migration-adjusted proliferation scores (Fig. 3c). Validation assays described above included examined variants in three residues in the GGG-binding pocket, Y104^3×33^I, K180^ECL2^I, and W181^ECL2^M, all of which showed decreased proliferation restraint and two of which showed decreased migration inhibition (Fig. 3g,i). Examination of the DMS results for all binding pocket residues showed significant intolerance to mutation for most positions (Fig. 5c). Positions with high expression-adjusted migration scores were significantly enriched among both the GGG-binding pocket positions and the Gα_13_-contacting positions (Fig. 5d). Similarly, 17/25 residues with high expression-adjusted migration scores were either part of the GGG-binding pocket or in contact with Gα_13_, a significant enrichment from 49/286 among non-CTD residues without high expression-ajusted migration scores (p < 0.0001, Fisher’s exact test).

**Fig. 5:**
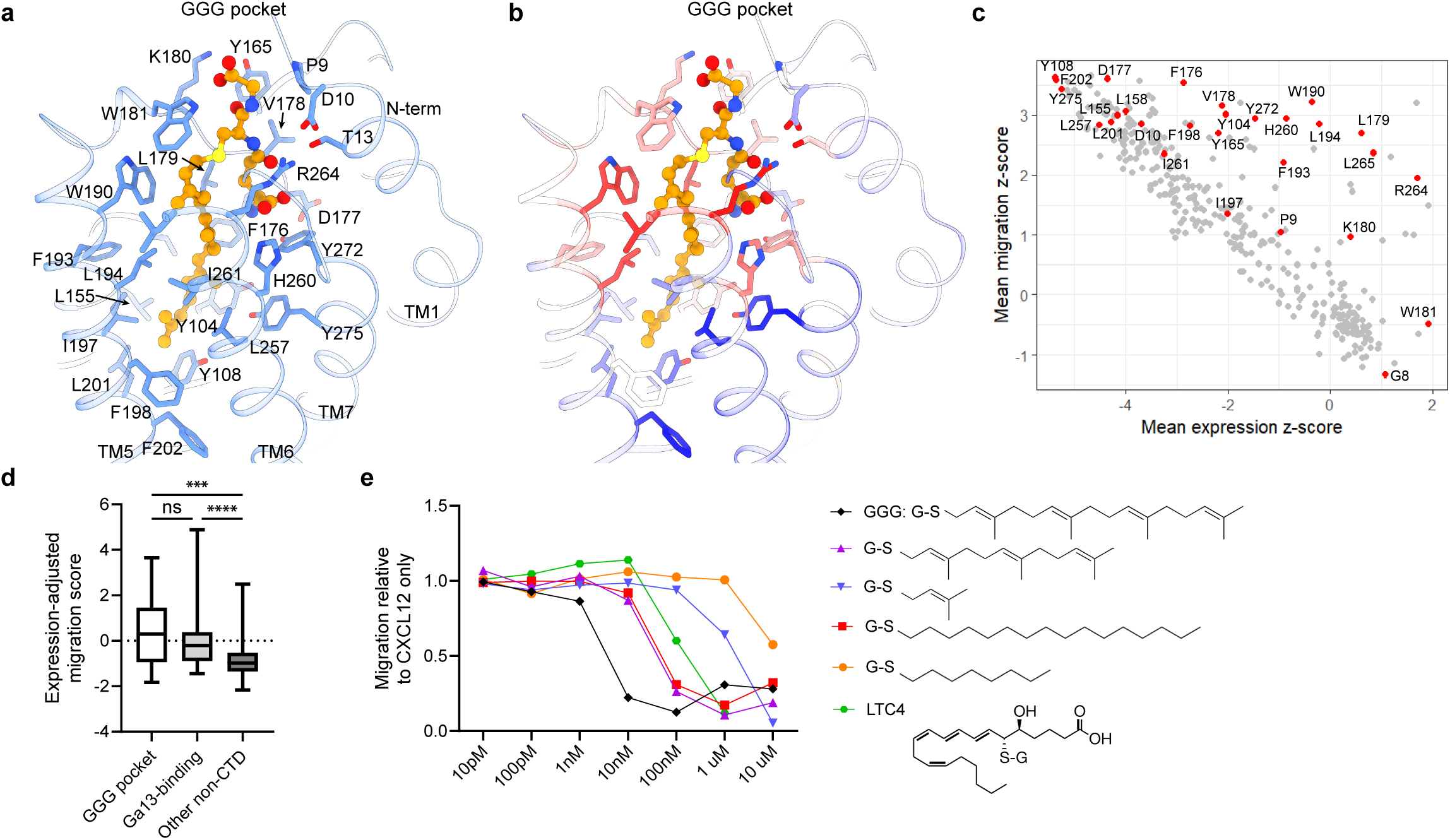
Structural features of GGG recognition. **a,b,** Close-up view of the GGG binding site in P2RY8. **A,** Expression effect sizes mapped onto P2RY8 residues (sticks and ribbon) that form the GGG binding site (orange sticks and spheres). **b,** Migration effect sizes mapped onto P2RY8 residues (sticks and ribbon) that form the GGG binding site (orange sticks and spheres). **c,** Plot comparing expression and migration mean z-scores by position, with GGG-interacting positions colored red. **d,** Distribution of expression-adjusted migration scores for GGG-binding pocket residues, Gα_13_-contacting residues, and other non-CTD residues. Whiskers extend to maximum and minimum, boxes shows 25^th^ percentile, median, and 75^th^ percentile. Kruskal-Wallis test with Dunn’s multiple comparisons test, ns = nonsignificant, *** < 0.001, **** < 0.0001. **e,** Assay of inhibition of P2RY8-transduced WEHI-231 cells, in the presence of 50 ng mL^−1^ CXCL12 with or without GGG or candidate P2RY8 ligands of varying concentrations, normalized to migration to CXCL12 alone. Representative results from one of two independent experiments. G-S, glutathione; LTC4, leukotriene C_4_.

Residues surrounding the deeper portion of the GGG-binding pocket were more tolerant to mutation (Fig. 5b). Our DMS results therefore suggest that the glutathione headgroup combined with the proximal prenyl group in the geranylgeranyl tail are required for P2RY8 activation. We tested our DMS-driven model for the most important chemical components of GGG required for P2RY8 activation with analogs. We previously showed that glutathione alone and geranylgeranyl-pyrophosphate are both inactive at P2RY8^29^. As we demonstrated previously the cysteinyl leukotriene C_4_ (LTC_4_) is active at P2RY8 with ∼100-fold less potency in a migration inhibition assay than GGG^29^. Although the specific orientation of LTC_4_ within the P2RY8 binding site is uncertain, it is likely that the glutathione headgroup and the aliphatic tail of LTC_4_ occupy similar general positions as GGG. The ability of LTC_4_ to activate P2RY8 suggested that the specific shape of the geranylgeranyl tail of GGG is not required for P2RY8 activation. Indeed, a saturated 16-carbon chain and a 15-carbon farnesyl group, which is one prenyl group smaller than geranylgeranyl, both activated P2RY8 with ∼10-fold less potency than GGG (Fig. 5e). Further diminutions in tail size significantly reduced potency, as seen for glutathione conjugated to a single prenyl group (∼1000-fold less potent than GGG) and glutathione conjugated to a saturated 8-carbon chain (∼5000-fold less potent than GGG). In sum, these results illustrated ligand potency varied with the degree of contact possible between the hydrophobic tail and P2RY8 within the binding site. The degree to which alterations in the glutathione head affect activity and potency remain an area for future research.

### Significant functional phenotypes with subtle proximal signaling changes

The specific points at which the P2RY8 signaling pathways affecting migration and proliferation diverge have not been established, but our screen results suggested that biased agonism of these activities was possible; on initial validation, V29^1×38^W (V29W) and W181^ECL2^M (W181M) showed WT-like behavior in migration but decreased proliferation restraint, whereas K180^ECL2^I (K180I) showed decreased but not absent function in both phenotypes (Fig. 3g,i). K180I and W181M are part of the GGG-binding pocket (Fig. 5a) whereas V29W is in TM1. We performed more in-depth validation of these variants to explore the mechanism for these phenotypes.

We first examined migration inhibition across a range of GGG concentrations, with results consistent with those from initial validation: W181M inhibited more strongly than WT at low GGG concentrations (3 nM) and comparably to WT at higher GGG concentrations; V29W was WT-like at all concentrations; K180I inhibited migration less than WT at all but very high GGG concentrations (Fig. 6a). A different pattern was observed when examining proliferation with a range of exogenous GGG concentrations. (We had not added exogenous GGG to the culture in the proliferation arm of the screen or initial validation because we had previously observed that B cells and B cell lines secrete GGG in amounts sufficient to affect proliferation^28,29,33^.) In this case, it was W181M that was most different from WT at lower GGG concentrations but equivalent at high concentrations; V29W had a similar but less striking pattern; K180I had a moderate deficit at all GGG concentrations relative to WT (Fig. 6b). In sum, although all three variants were able to restrain proliferation and migration when exposed to high levels of GGG, at low levels W181M showed greater defects in proliferation restraint and K180I greater defects in migration inhibition (Fig. 6a-c). This provided evidence that the P2RY8 signaling that drives these two processes is different at the receptor itself.

**Fig. 6:**
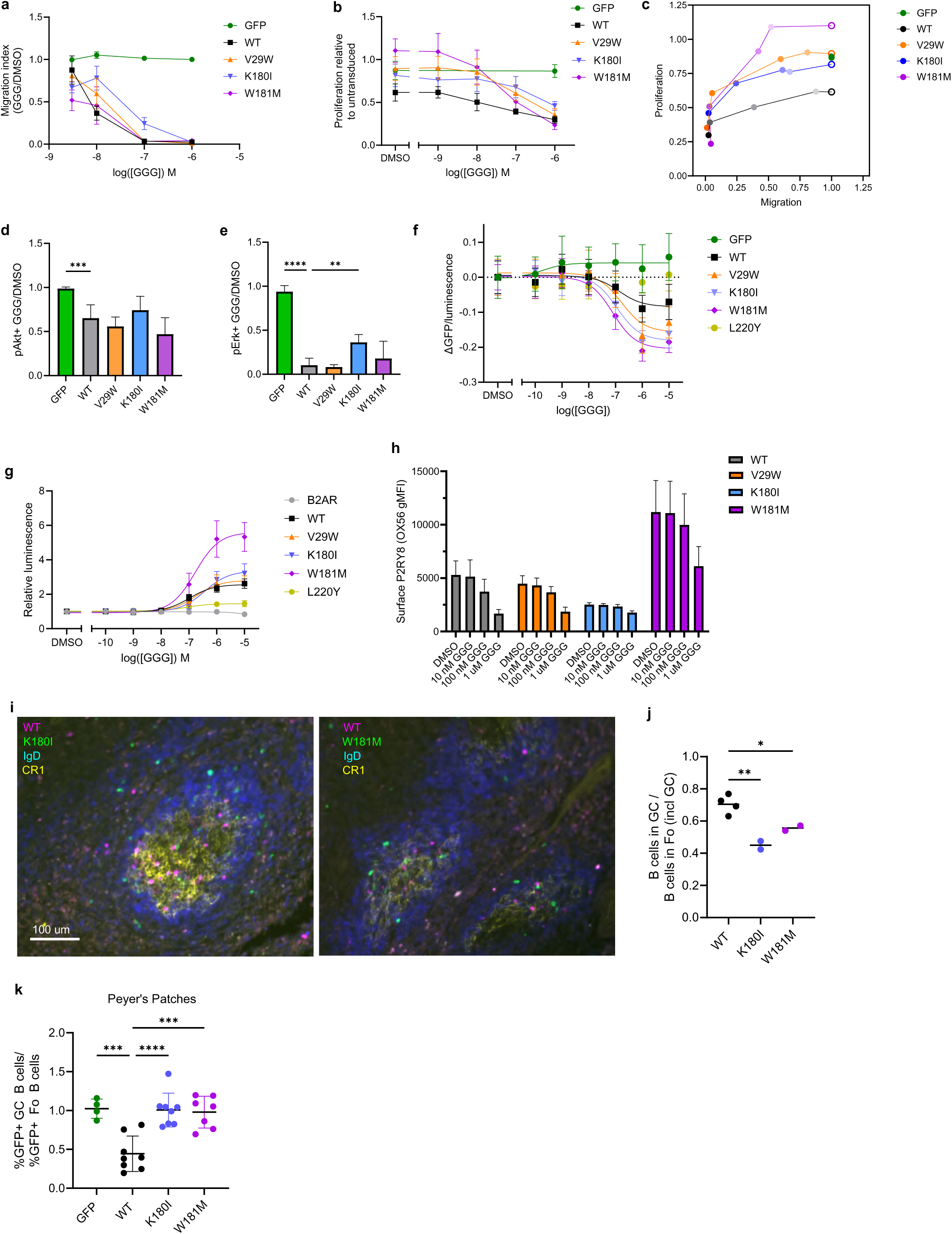
In vitro and in vivo delineation of heterogeneous variants effects. **a,** Migration of transduced Ly8 cells toward CXCL12 with varying concentrations of GGG, relative to DMSO (vehicle), and normalized to untransduced cells; points are means, error bars are SDs. Pooled results from 10 experiments, total 2 to 22 (mean 9.5, SD 4.9) replicates per condition. **b,** Proliferation over 7 days relative to untransduced cells in same well, with varying concentrations of exogenous GGG; points are means, error bars are SDs Pooled results from 5 experiments, total 6 to 12 replicates per condition. **c,** Migration and proliferation phenotypes at different GGG doses (comparing data from **a** and **b**). Empty circle indicates DMSO (vehicle), and ascendingly dark shading indicates 1 nM proliferation/3 nM migration, 10 nM, 100 nM, and 1 μM. **d,e** Transduced Ly8 cells exposed to GGG or vehicle were assayed using phospho-flow cytometry; graphs show ratio of pAkt^+^ (**d)** or pErk^+^ (**e)** cells in GGG vs vehicle, normalized to GFP (empty vector). Bars are means, error bars are SDs. Pooled results from 4 experiments, total 6 replicates per condition. **f,** BRET assay: 293T cells, co-transfected with P2RY8 variant, RLuc-Gα_13_ fusion, Gβ, and Gγ-GFP fusion, were treated with varying GGG concentrations and ratio of luminescence and GFP fluorescence determined, ratio normalized to DMSO (vehicle) condition. Points are means, error bars are SDs. Curves fit using log(inhibitor) vs response, 3 parameter model (Prism, Graphpad). Pooled results from 4 experiments, total 9 to 12 replicates per condition. **g,** NanoBiT assay: 293T cells co-transfected with GPCR-LgBiT and β-arrestin-SmBiT, treated with varying concentrations of GGG. Luminescence relative to DMSO (vehicle) plotted. Points are means, error bars are SDs. Curves fit using log(agonist) vs response, 3 parameter mode (Prism, Graphpad). Pooled results from 4 experiments, total 12 replicates per condition. **h,** P2RY8 surface expression of transduced Ly8 cells after 45 minute treatment with varying concentrations of GGG. Pooled from 3 experiments, total 6 to 8 replicates per condition. **i,** Representative immunofluorescence micrographs; activated polyclonal B cells transduced with K180I-GFP or W181M-GFP were co-transferred (∼3:2 ratio) with WT-TagBFP into SRBC immunized recipient mice; anti-GFP (green), anti-TagBFP (magenta), anti-CR1 (yellow, GC label), and IgD (blue, follicular B cell label); scale bar 100 μm, both images same scale. **j,** Quantitation of co-transferred transduced cells: ratio of transduced (GFP^+^ or BFP^+^) cells within the GC to those within the follicle (including GC), each dot is one mouse (a given mouse is represented by one WT dot and one K180I or W181M dot). Two independent experiments, one recipient mouse per variant per experiment, at least 45 GCs analyzed per mouse. **k,** Irradiated CD45.1 mice were reconstituted with bone marrow transduced with EV-GFP, WT-P2RY8-GFP, K180I-GFP, or W181M-GFP. After reconstitution, Peyer’s patches were analyzed for the frequency of GFP+ cells among GC and follicular B cells and the ratio plotted. Each point is a mouse. Mean and SD shown. Pooled from 3 experiments, total 4 to 8 mice per condition. **d, e, j, k,** One-way ANOVA test with Dunnett’s multiple comparisons test, each variant compared to WT; adjusted p-value ** < 0.01, *** <0.001, **** < 0.0001.

We then assayed elements of the known P2RY8 signaling pathway. It has previously been shown that P2RY8 activation results in decreased phosphorylation of Akt and Erk and that deleterious P2RY8 variants can disrupt this effect^29,33^. We used phospho-flow to assess simultaneously the phosphorylation status of these two kinases for the three variants of interest. As expected, WT P2RY8 allowed 100 nM GGG to trigger a significant reduction in the frequency of pAkt^+^ cells. Similar reductions were seen for all three interrogated variants (Fig. 6d). For all variants, GGG also drove a decrease in pErk^+^ cells, but K180I did so less effectively than WT (Fig. 6e). Overall, the phospho-flow results showed less separation between WT P2RY8 and the variants than was seen in the migration and proliferation assays.

We then used two luciferase-based reporter assays to assess variant effects on signaling pathway elements more proximal to the receptor. To examine trimeric G-protein activation, we used TRUPATH, a BRET-based approach^42,43^. In this system, luciferase is fused to Gα_13_; when it encounters its substrate, it can trigger fluorescence of GFP fused to Gγ. Activation of the trimeric G-protein results in dissociation of the complex and decreased GFP signal relative to luciferase signal. L220^5×65^Y, a residue that interacts with Gα_13_ and was deleterious in initial validation studies (Fig. 3g,i), was unable to drive G-protein activation (Fig. 6f). In contrast, WT P2RY8, V29W, K180I, and W181M were all able to do so in a dose-responsive manner with comparable IC_50_ values (Fig. 6f). Although the variants showed a larger absolute change than WT, the degree to which this reflected differences in expression versus intrinsic per-receptor maximum activity was not determined. To measure β-arrestin recruitment, we used the NanoBiT system^44^. In this system, a large fragment of luciferase is fused to the GPCR and a small fragment to β-arrestin. When the GPCR and β-arrestin are in close proximity, the two luciferase fragments interact to form a functional luciferase. As expected, a GPCR (β2AR) with a different ligand from P2RY8 was unable to recruit β arrestin with the addition of GGG (Fig. 6g). In contrast, WT, V29W, K180I, and W181M all showed robust recruitment, with W181M having higher maximum recruitment, though not significantly different EC_50_ (Fig. 6g). L220^5×65^Y showed significantly reduced recruitment but was distinguishable from β AR. In sum, neither of these assays provided a mechanistic explanation for the differential effects of these variants on migration and proliferation.

For many though not all GPCRs, β-arrestin recruitment promotes receptor internalization^19^. We examined changes in receptor surface expression after 45 minutes of exposure to various concentrations of GGG. V29W showed a similar pattern to WT, whereas W181M had a higher baseline expression and less relative decrease at 100 nM, and K180I had a lower baseline expression but only modest reduction in surface expression even at high GGG concentrations (Fig. 6h). The results of the NanoBiT β-arrestin assay and this internalization assay together suggest that the ability to recruit β-arrestin is not sufficient to produce a normal P2RY8 internalization response to ligand. In sum, although dose-response studies further validated the migration and proliferation phenotypes, assays of proximal signaling elements did not delineate the specific mechanism underlying these phenotypes.

### Disparate migration and proliferation phenotypes of two P2RY8 variants in vivo

Given their contrasting phenotypes in vitro as described above (Fig. 6a-c), we decided to further assay the function of K180I and W181M in vivo. P2RY8, although conserved across vertebrates, has been lost in rodents. However, previous work has illustrated that human P2RY8 introduced into mice is able to function to affect B cell positioning in lymphoid follicles and GC B cell abundance in Peyer’s patches^28,29^. To examine the effect of these mutants on B cell positioning in lymphoid tissues, polyclonal murine B cells were activated in vitro, transduced with variant or WT P2RY8, and then co-transferred into pre-immunized mice before harvest for immunofluorescent microscopy analysis. The variant vectors contained GFP whereas the WT vector contained TagBFP, permitting direct comparison within the same recipient mice. We have previously shown that in the absence of P2RY8, transferred activated B cells do not localize into the GC, whereas most cells with WT P2RY8 congregate within the GC^28^. We observed this same localization pattern with WT P2RY8-transduced B cells, with approximately 80% of TagBFP^+^ cells within the follicle being located in the GC (Fig. 6i,j). For both K180I and W181M, approximately half of the transduced cells within the follicle were in the GC (Fig. 6i,j). This in vivo phenotype is consistent with the modest migration inhibition defect seen in in vitro for K180I. The mechanism for the decreased localization of W181M in vivo compared to WT, despite at least WT-like responsiveness in vitro, is less evident. Although previous direct GGG measurement in lymphoid tissue homogenates found a concentration of approximately 10 nM^29^, the interstitial concentrations at different sites within the lymphoid tissue are uncertain aside from a relatively lower level within the GC^28,29^. The in vivo behavior differences for W181M compared to WT that contrast with the in vitro migration assay (Fig. 6a) may be a consequence of other differences resulting from this variant, including higher baseline surface expression and decreased internalization in response to ligand, differences in response kinetics, and more complicated chemokine and GGG gradients in vivo compared to the transwell assay.

In a second in vivo assay, we used transduced bone marrow to reconstitute irradiated congenic mice, and after reconstitution we compared the proportion of transduced cells among GC and follicular B cells in three tissues. We have previously shown that WT P2RY8-transduced cells are less prevalent among GC B cells than follicular B cells in Peyer’s patches, whereas there is not a significant difference in mesenteric lymph nodes or spleen^28^. The current experiments reproduced this effect for WT; in contrast, neither K180I nor W181M showed altered prevalence in GC compared to follicular B cells in any of the three tissues (Fig. 6k and Extended Data Fig. 9a,b). This argues that the phenotype produced by WT P2RY8 requires a high degree of receptor activity, and that even moderate defects in proliferation restraint, as seen in vitro with K180I (Fig. 5b), result in inability to recapitulate the WT in vivo growth-regulation phenotype. The still greater deficit of W181M at physiologic ranges of GGG in vitro (Fig. 5b) would therefore also translate to inability to recapitulate the phenotype produced by WT P2RY8. Overall, the results of these in vivo assays provide the first evidence that it is possible for P2RY8-driven confinement within the GC to occur without causing decreased prevalence within the GC B cell pool.

### Phenotypes of germline and lymphoma-associated P2RY8 variants

Hundreds of missense variants in P2RY8 have been observed in human population sequencing studies, distributed across the protein and with highly varied phenotypes per our DMS results (Fig. 7a). Most such variants are rare or very rare but comparison of allele frequency in the gnomAD data set^2^ with their DMS effect sizes nonetheless revealed statistically significant correlations, consistent with selective pressure. These are apparent both when the 476 individual variant frequencies are considered and when consolidating these variants within the 260 involved positions (Fig. 7b and Extended Data Fig. 10a-e). Four of the variants validated above (Fig. 3e,g,i) are among the variants reported in gnomAD (F103^3×32^L, which shows LoF, and G237^6×36^N, A240^6×39^T, and T330^CTD^N, which are WT-like).

**Fig. 7:**
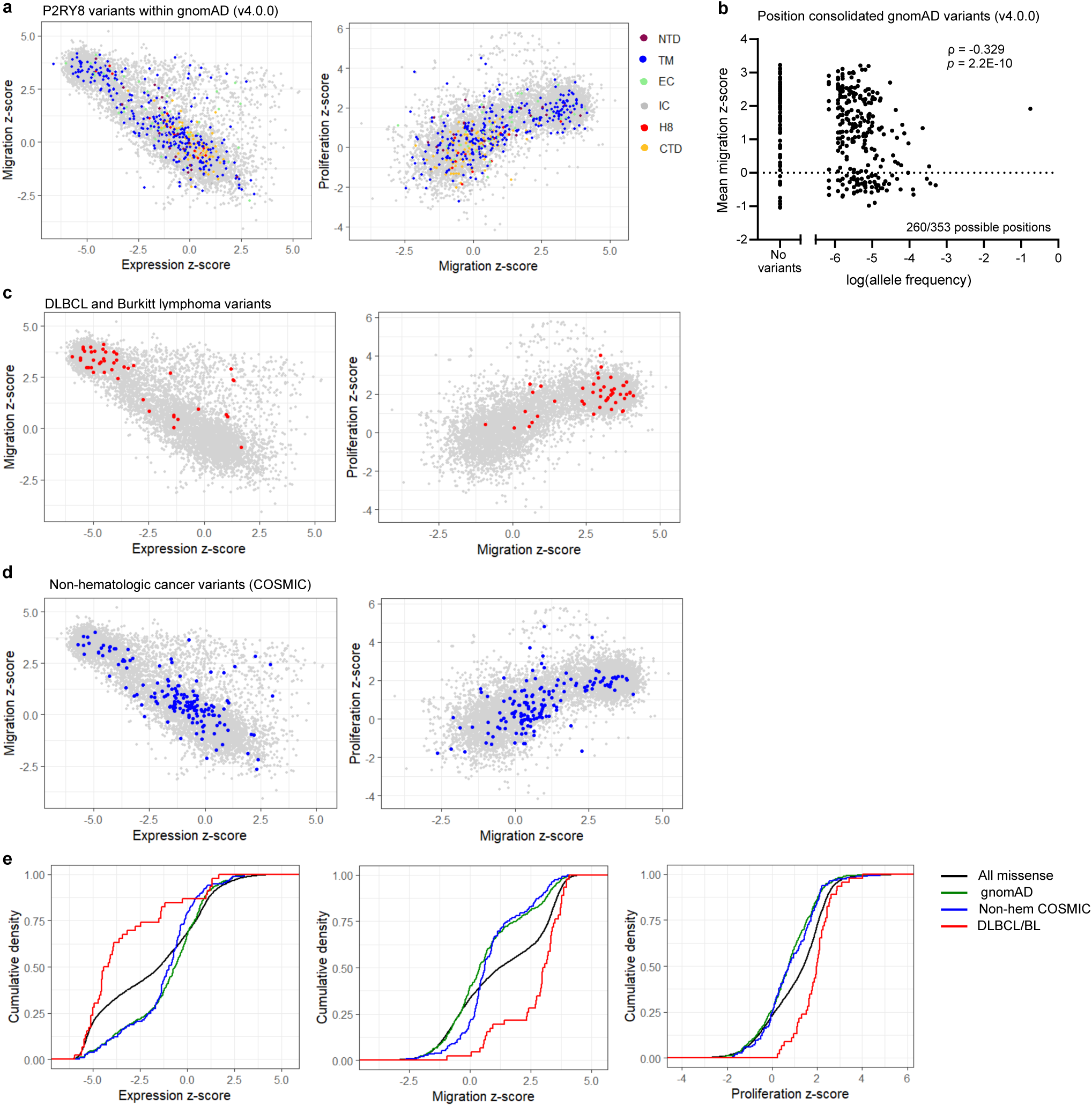
Phenotypes of germline and lymphoma-associated P2RY8 variants. **a,** Plots comparing expression and migration z-scores (L) and migration and proliferation z-scores (R) of variants, with human population data in gnomAD (v4.0.0) colored by domain of protein. **b,** Plot of allele frequency for missense variants in gnomAD (v4.0.0), summed within each position, to the mean migration z-score for each position. ρ, Spearman correlation; p-value calculated using algorithm AS 89 with Edgeworth series approximation. **c,** Plots comparing expression and migration z-scores (L) and migration and proliferation z-scores (R) of missense variants, with 48 variants reported in DLBCL or Burkitt lymphoma colored red. **d,** Plots comparing expression and migration z-scores (L) and migration and proliferation z-scores (R) of missense variants, with non-hematologic cancer variants colored blue **e,** Cumulative density plots of expression, migration, and proliferation z-scores for all missense variants (black), gnomAD (v4.0.0) variants (green), non-hematologic cancer variants (blue), and DLBCL / Burkitt lymphoma variants (red).

P2RY8 missense variants are common in two GC B-cell-derived lymphomas, DLBCL and Burkitt lymphoma. We plotted the effect sizes of 48 variants compiled from two sources (COSMIC^32,45^ and Muppidi et al.^26^) and observed that nearly all of these variants reduced P2RY8 expression and decreased its migration inhibition, and none of them showed increased expression or function relative to WT (Fig. 7c). Although 15 of these variants have been observed in gnomAD as germline variants, none have an allele frequency > 0.0001, indicating it is very likely that they were somatically acquired in these cancers. Of the 41 positions with lymphoma variants, six were identified above (Fig. 3a) as having high expression-adjusted migration scores (F103^3×32^, Y104^3×33^, A191^5×36^, A235^6×34^, R264^ECL3^, and F266^ECL3^). In addition, another involved position, Y30^1×39^, has the highest mean proliferation effect size of any position (Fig. 3c). These lymphoma variants involve residues found to interact with Gα_13_, including M62^2×43^, A235^6×34^, and S297^8×47^, and residues in the GGG-binding pocket, including Y104^3×33^, F176^ECL2^, D177^ECL2^ and R264^ECL3^. Of the three lymphoma variants with a large migration effect size but no decrease in expression (Fig. 7c), two involve R264^ECL3^: R264C and R264H. In addition, our screen revealed that six of the lymphoma variants that are LoF would not have been classified as pathogenic based on AM pathogenicity score (P20^1×29^L, R86^ECL1^C, A140^4×42^T, R264^ECL3^H, S270^ECL3^N, and K276^7×36^R; Supplemental Tables 1 and 2).

Turning to 159 missense variants that have been identified in non-hematologic cancers (COSMIC), we observed a broadly distributed set of phenotypes akin to that seen in germline variants (Fig. 7d). Although 67 of these variants are present in gnomAD, only 3 have an observed allele frequency > 0.0001 (V125^3×54^I, R133^34×55^P, V333^CTD^L), again suggesting predominantly somatic mutagenesis. Ten of these variants are also present in the DLBCL/Burkitt lymphoma set. Comparison of the phenotype distributions for these four sets of variants using Kolmogorov-Smirnov tests revealed statistically significant differences, with gnomAD and non-hematologic cancer variants containing fewer deleterious variants than the set of all missense variants, and DLBCL/Burkitt lymphoma variants being enriched for deleterious variants (Fig. 7e and Extended Data Fig. 10f). These findings aligned with previous observations supporting P2RY8 as a cancer driver gene in DLBCL and Burkitt lymphoma, in contrast to non-hematologic cancers in which P2RY8 variants are more likely passenger mutations.

## Discussion

Here, we report near-saturation DMS of human P2RY8, a G_13_-coupled GPCR important for GC B cell confinement, defining the expression, migration, and proliferation phenotypes for these variants. Use of these different phenotypic readouts allowed for delineation of variants that affected both expression and function, affected function independently of expression, and discrepantly affected migration and proliferation.

VEP algorithms have improved significantly and, particularly at the position level, are highly accurate at identifying changes that will result in loss of protein function. In P2RY8, however, these methods are much less able to predict increases in protein expression or activity, echoing similar limitations in VEP analysis of other proteins^7,9^. Our work demonstrates that it is possible to generate new, more highly correlated prediction scores through the combination of VEP scores and limited amounts of experimental data; even 2 variants per position resulted in a substantial improvement. Although greater gains were achievable with actual fine-tuning of the protein language model itself, even an OHE regression-based approach, which does not require knowledge of the model weights, achieved significant improvements. Despite these sizeable improvements in predictions for P2RY8 itself, use of P2RY8 experimental data could only modestly improve variant effect predictions for a recently profiled GPCR, GPR68. It is possible that alternative approaches to fine-tuning might enhance the ability to generalize from one receptor to another. Our results suggest that predictions across a family of GPCRs can be achieved more efficiently by profiling relatively few variants across many receptors, rather than comprehensively profiling variants within a small number of receptors. Beyond single missense variants, future studies could assay the combinatorial effects of multiple variants, indels, and other structural variants, in order to enhance our understanding and ability to computationally predict epistasis in GPCRs.

The structure of activated P2RY8 provides further understanding of the functional data obtained through the DMS. By identifying key residues involved in interactions with Gα_13_ and GGG, as well as our work evaluating the potency of several alternative ligands, we lay the groundwork for additional studies aimed at developing alternative agonist and antagonist ligands of this receptor.

Our exploratory studies of in-depth validation of select variants highlighted areas for continued investigation. In particular, the mechanism by which a given variant may produce different degrees of defects in migration or proliferation restraint remains unclear, particularly as the assays of trimeric G-protein activity and β-arrestin recruitment did not show clear differences that might explain these phenotypes. This highlights the ability of our phenotypic screens to detect subtle phenotypes that may otherwise be missed in classical GPCR signaling assays that report changes more proximal to the receptor. We believe that results from our screens provide a more complete picture of how P2RY8 activation manifests as a phenotype, by intergrating the compounding and amplified effects that resulting in subtle shifts in signal propagation through the cell. Future study into the basis for these variants’ phenotypes will be useful in further defining P2RY8 function. Additionally, P2RY8 variants identified in this study will enable in vivo studies to probe the role of P2RY8 function in GCs^29^, B cell negative selection^33^, and T cell responses^28^.

One caveat of our work is that the range of observed effect sizes is greater for expression than for migration and both are greater than for proliferation. This may reflect details of the screen itself, in which the maximum possible enrichment differs between the readouts. Adjustments, such as running the proliferation assay for a longer period of time, could conceivably result in increased effect sizes thereby enhancing phenotyping of proliferation.

Our work provides a resource for interpretation of the pathogenicity of nearly any given missense variant of P2RY8, whether germline or somatic, which may be of particularly utility in the setting of autoimmunity where deleterious P2RY8 variants have already been identified in a small number of patients^33^. It also allows interpretation of any somatic P2RY8 variants observed in cancer, including identification of deleterious variants that may not be identified by current computational VEP tools. Using transfer learning, our results may also be used to facilitate the analysis of other GPCRs.

In conclusion, we have used DMS and cryo-EM to comprehensively map the mutational landscape of the immunomodulatory GPCR P2RY8, advancing understanding of its function.

## Methods

### Cell lines

HEK 293T (293T), WEHI-231, and parent OCI-Ly8 (Ly8) cell lines were previously obtained from other laboratories and further authentication was not performed. The cell lines were not tested for Mycoplasma contamination. P2RY8 KO Ly8 cells were previously generated as described^29^. The culture medium for Ly8 and WEHI-231 cells was RPMI-1640, 10% FBS, 1x GlutaMax, 10 mM HEPES, 55 uM β-mercaptoethanol, and 50 IU penicillin/streptomycin. Lenti-X 293T cells (Takara) were used to generate lentivirus for transduction of human cells. The culture medium for Lenti-X was DMEM, 10% FBS, 1x GlutaMax, 1 mM sodium pyruvate, 1x MEM non-essential amino acids, 10 mM HEPES, and 50 IU penicillin/streptomycin. Plat-E 293T (Plat-E) cells (Cell Biolabs) were used to generate retrovirus for transduction of mouse cells. The culture medium for Plat-E and 293T cells was DMEM, 10% FBS, 1x GlutaMax, 10 mM HEPES, and 50 IU penicillin/streptomycin. All cells were maintained in a 37°C humidified incubator, 5% CO_2_.

### Mice

Mice used for B cell transfers were C57BL/6J bred in an internal colony and used at 8 to 12 weeks of age. For chimeras, donor bone marrow was obtained from C57BL/6J bred internally or purchased from JAX; recipients were CD45.1 B6 (B6.SJL-*Ptprc*a*Pepc*b/BoyCrCrl) mice, bred internally from founders ordered from JAX or purchased from the National Cancer Institute at Charles River at age 7 to 8 weeks, with balanced distribution of experimental groups across these two backgrounds. Mice were allocated to control and experimental groups randomly, and sample sizes were chosen based on previous experience and available co-caged littermates. Animals were housed in a pathogen-free environment in the Laboratory Animal Resource Center at UCSF, and all experiments adhered to ethical principles and guidelines that were approved by the Institutional Animal Care and Use Committee.

### Variant pool generation

The vector for library transduction was created through modification of p_sc-eVIP (p_sc-eVIP was a gift from Jesse Boehm, James T. Neal, and Aviv Regev; Addgene plasmid # 168174)^46^. First we removed the puromycin resistance gene by digestion with XhoI, gel purification (Macherey-Nagel Nucleospin kit), blunt end formation withT4 DNA polymerase (NEB), and ligation with Quick Ligase (NEB), transformation into chemically competent bacteria, and plasmid isolation and screening. We then excised P53 and replaced it with PRL-OX56-P2RY8-P2A-GFP. Specifically, two separate PCR reactions were performed using Q5 polymerase (NEB) on MSCV-PRL-OX56-P2RY8-IRES-GFP plasmid (previously described^29^) to produce PRL-OX56-P2RY8-P2A and P2A-GFP fragments with appropriate homology arms for Gibson assembly and thereby cloned into BmtI and PmeI digested plasmid backbone using Gibson assembly kit (NEB). This resulting plasmid is termed POP2E.

The variant oligo library was obtained as Site Saturation Variant Library from Twist. The synthesized DNA consisted of the coding sequence for P2RY8 amino acids 2-358 (all codons excepting start and stop codons) along with preceding 35 bp (corresponding to OX56 tag and 5 nucleotides in preceding linker) and succeeding 35 bp (corresponding to a portion of P2A sequence). The pools was designed to include all possible missense mutations (a single codon for a given missense amino acid if multiple codons were possible), along with a synonymous codon for all amino acids for which multiple codons exist (that is, all except methionine and tryptophan). Site-directed mutagenesis with the Q5 site-directed mutagenesis kit (NEB) was to modify POP2E by inserting a CTA between P2RY8 and P2A, producing a AvrII restriction site. This new construct was digested with PshAI and AvrII, the non-P2RY8 fragment isolated by gel cleanup (Qiagen Qiaquick Gel Extraction kit), and the oligo pool inserted using using HiFi Assembly kit (NEB): 0.0625 pmol variant oligo library and 0.25 pmol of backbone were combined with 2x master mix for 60 minutes at 50°C. The reaction was cooled on ice then dialyzed for 1 hour on a 0.025 μm MCE membrane (Millipore) floating on ultra-distilled water. Added 2 μL of dialyzed assembly reaction to 50 uL of Endura electrocompetent cells (Biosearch Technologies), incubated on ice for 15 minutes, aliquoted equally to two 0.1 cm cuvettes (Bio-Rad), and electroporated with 1.8 kV, 10 uF, 600 Ω pulse. This was combined with pre-warmed SOC medium, shaken at 37°C for 1 hour, and spread across two 25 cm x 25 cm LB ampicillin plate, with the exception of a small volume used in serial dilutions to determine transformation efficiency. Based on these serial dilutions, the large plates had a total of 1.67 × 10^6^ transformants, for an average of 233 transformants per variant. After overnight growth, colonies were scraped from the plates, pooled, divided in six, processed using HiSpeed Plasmid Midi Prep kit (Qiagen), and repooled.

To produce lentivirus, Lenti-X cells were plated into two 10 cm plates, with transfection the following day with cells ∼85% confluent. To 3 mL of Opti-MEM, added 90 uL of lipofectamine 3000; to another 3 mL Opti-MEM added 13 μg of variant library plasmid, 14 μg psPAX2, 3.5 μg pMD2.G, and 80 μL of P3000 reagent. Combined these after 5 min room temperature incubation. After 25 min further incubation at room temperature, removed culture medium from the Lenti-X plates, then added 5 mL of complete Opti-MEM (Opti-MEM with, 5% FBS, 1x GlutaMax, 1 mM sodium pyruvate, and 1x MEM non-essential amino acids) and 3 mL of the lipofectamine-DNA mixture to each plate. Six hours later removed media and replaced with 6 mL of complete Opti-MEM plus 50 IU penicillin/streptomycin and 1:500 ViralBoost (Alstem). After 24 hours, the media was collected, passed through a 0.45 μm filter, and concentrated 20x using Lenti-X concentrator (Takara) and centrifugation. Four such batches of lentivirus were produced for use in the screen.

### Variant pool screening

The screen was performed using Ly8 cells in which P2RY8 knockout by CRISPR-Cas9 had previously been performed^29^. For each iteration of the screen, 7 × 10^6^ cells were transduced by resuspending cells at 1 × 10^6^ cells mL^−1^ in culture medium and 750 μL of lentivirus diluted in Opti-MEM, such that ∼30% of cells were transduced (range 25-34%), indicating an average of 294 transduced cells per variant. Post-transduction, the cells was passaged as needed to maintain a cell concentration of 1.5 × 10^5^ to 1.5 × 10^6^ cells mL^−1^ with a total of at least 3 × 10^6^ transduced (GFP^+^) cells. The four replicates for each assay (surface expression, migration, and proliferation) were drawn from seven independent transductions using four batches of lentiviral library.

Sample collection for surface expression assay occurred 5 days after transduction for 3 replicates and 7 days after transduction for 1 replicate. Cells were washed with FACS buffer (1x PBS, 2% FBS, 1 mM EDTA), stained for 30 minutes on ice with 1:100 TruStain FcX block (Biolegend) and 1:200 biotinylated anti-OX56 (Bio-X-Cell) at 40 × 10^6^ cells mL^−1^, washed with FACS buffer, stained for 30 minutes on ice with 1:400 streptavidin-AF647 (Fisher), with e780 fixable viability dye (eBioscience) added to 1:1500 for last 10 minues, washed with FACS buffer, resuspended in FACS buffer, and sorted using FACSAriaFusion. Gating was FSC-A x SSC-A, singlets by FSC-A x FSC-W, live cells by FSC-A x e780, GFP+, and four bins of OX56 (AF647) expression, containing 20%, 30%, 30%, 20% (Extended Data Fig. 1b). Cells were sorted into 25% FBS in PBS at 4°C. Cells were washed, pelleted, and frozen at −80°C until DNA extraction for processing and sequencing. An average of 2 × 10^6^ cells per bin per sort were collected.

The collection for proliferation analysis occurred at 3, 5, and12 days after transduction for all four replicates asas well as 7 days after for two replicates and 8 days after for one replicate. (The Enrich2 algorithm can accommodate and account for these differences.) The proportion of GFP^+^ cells was determined by flow cytometry of a small aliquot at the time of collection. Cell aliquots containing an average of 2.3 × 10^6^ GFP^+^ cells (range 1.5-3.1 × 10^6^) were washed with PBS, pelleted, and frozen at −80°C until DNA extraction for processing and sequencing.

Migration assays occurred 5 days (one replicate), 7 days (two replicates), or 8 days (one replicate) after transduction. In each case, 80 × 10^6^ cells were washed with migration medium (RPMI, 0.5% fatty acid-free bovine serum albumin, 1x penicillin/streptomycin), resuspended in migration medium at 1 × 10^7^ cells mL^−1^ and resensitized for 15 minutes at 37°C. Migration medium containing 100 ng mL^−1^ recombinant human CXCL12 (Peprotech) and 100 nM GGG (Cayman) was prepared and 600 μL added to each of 80 wells in 24-well tissue culture plates. Transwell inserts (6 mm, 5 uM pore size, Corning) were placed in each well, after which 100 uL resensitized cells (1 × 10^6^ cells) were added to the upper chamber. The cells were placed in 37°C, 5% CO_2_ incubator for 3 hours to allow migration. The transwells were then removed and the cells in the bottom wells were pooled and counted. A small aliquot was also analyzed by flow cytometry to determine the proportion of GFP^+^ cells. The total number of migrated cells ranged from 10 to 21 × 10^6^ cells. In addition, an aliquot containing 2-3 × 10^6^ GFP^+^ cells from the pre-migration input population was also separately collected for each migration. Cells were washed with PBS, pelleted, and frozen at −80°C until DNA extraction for processing and sequencing.

### Sequencing library preparation

Library preparation and sequencing was performed as two batches, with two replicates for each condition per batch. Genomic DNA was isolated using Quick-DNA Miniprep Plus kit (Zymo) per its instructions. DNA was quantified using Qubit (Thermo Fisher Scientific). Initial PCR amplification reactions were set up to amplify P2RY8 coding sequence plus ∼150 flanking bp on each end. Each reaction was 50 μL using Q5 High-Fidelity Polymerase (NEB), including 0.5 μL polymerase, 10 μL 5x buffer, 10 μL high GC enhancer, 0.5 μL 20 μM forward primer, 0.5 μL 20 μM reverse primer, and 27.5 μL of DNA, up to 1500 ng of DNA per reaction. When possible, number of individual PCR reactions per sample was set to have total template equivalent to 150 transduced cells per variant (1.057 × 10^6^ transduced cells total); achieved for 26 of 39 samples; variant coverage for the remaining samples was 57x, 66x, 83x, 100x (four samples), 104x, 106x, 120x, 124x, 145x, and 146x. PCR conditions were 98°C x 2 min, 19 cycles of (98°C × 10 sec, 70°C x 15 sec, 72°C x 45 sec), and 72°C x 2 min. After the PCR was performed, all individual PCR reactions for a given template were pooled. Approximately 5-10% (depending on total number of reactions) of each pooled reaction was loaded onto agarose gel, band of desired size verified using gel electrophoresis, and band then cut out and PCR product isolated using Qiaex II kit (Qiagen). DNA was quantified using Qubit. DNA was diluted as needed and 1 ng was tagmented using Nextera XT Library Prep kit (Illumina) per its instruction, including 12 cycle PCR for adding Illumina adapter sequences. DNA clean up performed using AMPure XP beads (Beckman) using 0.75x volume (to remove fragments <300 bp). Quantification and QC was performed using Qubit and gel electrophoresis including TapeStation high-sensitivity electrophoresis (Agilent). Samples were diluted and pooled in equimolar fashion to 10 nM total. PE300 sequencing was performed on Illumina NovaSeq X Plus at UCSF CAT.

### Sequencing analysis

Fastq files were obtained from UCSF CAT. There was a mean of 6.51 × 10^7^ paired sequences per sample, SD 1.02 × 10^7^, max 8.60 × 10^7^, min 3.72 × 10^7^. Overall quality was evaluated using FastQC^47^ (version 0.12.0) (http://www.bioinformatics.babraham.ac.uk/projects/fastqc/). Adapter trimming then performed using BBDuk function of BBTools^48,49^ (version 38.18) (https://sourceforge.net/projects/bbmap/). Paired reads were error corrected using BBMerge (also part of BBTools). Reads were then mapped to P2RY8 amplicon reference sequence using BBMap (also part of BBTools). Variants were then called in the mapped SAM file using AnalyzeSaturationMutagenesis function within GATK^50,51^ (version 4.5.0.0) (https://gatk.broadinstitute.org/hc/en-us). The utilized output of this analysis was a table containing the read count for each codon for each position in the coding sequence of P2RY8. These tables were then processed using R to filter for only the codons specifically used by Twist in creating the variant oligo pool and formatted as required for analysis by Enrich2^52^ (version 1.3.1)(https://github.com/FowlerLab/Enrich2/). Enrichment scores were calculated from the processed files, looking at all four replicates concurrently, by weighted least squares method for surface expression and proliferation and log ratios for migration (since only two points). For each sample, the geometric mean of the counts for the 334 synonymous variants had been calculated and was used as the sample WT count for normalization within Enrich2. The Enrich2 output was further processed in R, including determination of z-scores for expression, migration, and proliferation for all missense variants based on the standard deviation of the Enrich2 scores for the synonymous variants. Expression-adjusted migration score determined by finding sum of expression and migration mean z-scores for each position, with high expression-adjusted migration score defined as being more than 1.5 times the interquartile-range away the median and having mean migration z-score > 0.75. Variant counts and Enrich2 scores are in Supplemental Table 2. Scripts used for analysis are available on GitHub at https://github.com/yelabucsf/P2RY8_DMS.

### ESM1b fine-tuning using P2RY8 DMS data

ESM1b takes a single amino acid sequence of length L as input and outputs a 20 x L matrix of log-likelihood ratios (LLR) corresponding to the predicted effects of all single amino acid substitutions^6,53^. To improve the predictive power of ESM1b on P2RY8, here we fine tune the model on a randomly chosen subset of the DMS data (e.g., expression z-scores of k variants per position) and use the rest of the DMS data for performance evaluation (test set). This process is repeated N=50 times for each experimental measurement (expression, migration and proliferation) and each training set size with k = 2, 3, 5, 7, 9 and 11 variants per position.

Fine-tuning was performed by updating the parameters of the ESM1b model to minimize a masked mean-squared error loss computed between the sampled experimental values and the corresponding model predictions. Before calculating the loss, the sampled experimental values were transformed to match the mean, variance, and the sign of the P2RY8 zero-shot LLRs to enable faster convergence and avoid major changes to the language model logit distribution during training. We further restricted optimization to the language model’s final layers, comprising the LM head and the last three embedding layers (∼60M trainable parameters), to preserve the broader contextual knowledge acquired during self-supervised pre-training while enabling adaptation to the experimental data.

Model optimization was carried out using an AdamW optimizer (with a learning rate of 1e-5 and weight decay of 0.01) alongside a linear warmup schedule for a maximum of 300 optimization steps. To avoid overfitting to the sampled experimental data, we implemented an early stopping strategy. Specifically, for each training set size, we fine-tuned the model on k-1 variants per position and used the remaining one variant per position for validation. Throughout training, the validation set was used to monitor model performance (Spearman correlation between model predictions and experimental measurements), and training was stopped if the validation correlation did not improve over a predefined number of steps (patience = 30 steps).

After fine-tuning, the updated model generated revised LLR predictions for all P2RY8 variants. To further refine these predictions, we incorporated a ridge regression step that combined one-hot encoded sequence features with the fine-tuned model scores. This final regression model was trained on the same set of k experimental variants per position (including variants from both training and validation sets) without transforming the experimental values, to provide predictions on the experimentally measured scale (e.g., P2RY8 expression z-scores in the case of the expression DMS screen). We also implemented the augmented one-hot encoded ridge regression model previously described^35^, which relies solely on zero-shot model predictions (AlphaMissense and original ESM1b scores), as a baseline to evaluate the performance gains obtained by fine-tuning ESM1b using P2RY8 DMS data.

Finally, we tested whether ESM1b fine-tuning on P2RY8 DMS data can improve variant effect predictions for GPR68, a related GPCR with available DMS data. For this task, we fine-tuned ESM1b on P2RY8 as above, using all experimental DMS data (expression z-scores) for training and tested performance on GPR68 (surface expression and ph 5.5 signaling). We tested the performance of two fine-tuned models, one solely relying on the P2RY8 validation set for early stopping and a second one with a hard stop at 50 optimization steps to avoid overfitting. Overall both models generate similar variant effect scores (LLR) when tested on GPR68, with the latter providing a small increase in Spearman correlation compared to the original (zero-shot) ESM1b model.

### Individual variant validation

Specific variants of interest were produced in the POP2E (WT P2RY8) vector by site-directed mutagenesis (Q5 Site-Directed Mutagenesis Kit, NEB). Lentivirus was produced using Lenti-X by same approach as for lentiviral pool (described above) but scaled down to 1-2 wells in a 6-well plate. Ly8 cells were transduced by same approach as lentiviral pool, again scaled down to 1 × 10^5^ cells in 100 uL in a well in a 96-well plate per transduction.

For surface expression, cells were assessed 3 days after transduction. Cells were washed with FACS buffer, stained for 30 minutes on ice with 1:100 TruStain FcX block (Biolegend) and 1:200 biotinylated anti-OX56 (Bio-X-Cell), washed twice, stained for 30 minutes on ice in the dark with 1:250 streptavidin-AF647 (Fisher), washed, stained for 10 minutes on ice in dark with 1:1500 e780 fixable viability dye (eBiosciences), washed, and resuspended and analyzed on a FACSymphony (BD). An aliquot of untransduced Ly8 cells were used in each experiment to set the GFP^+^ gate.

For proliferation, transduced cells were maintained in wells of a 24-well tissue culture plate. The ratio of GFP^+^ to GFP^−^ cells was tracked over time by performing flow cytometric analysis of an aliquot at 3, 5, 7, 10, 13, and 16 days after transduction. Cells were passaged at those same time points.

For migration, cells were assayed 5-7 days after transduction. Cells were washed with migration medium (RPMI, 0.5% fatty acid free BSA (Sigma), 10 mM HEPES, and 50 IU penicillin/streptomycin), resuspended at 2 × 10^6^ cells mL^−1^ and resensitized for 10 to 15 minutes at 37°C. Recombinant human CXCL12 (Peprotech) was diluted to 100 ng mL^−1^ in migration medium. GGG was diluted to various concentrations in the CXCL12-containing migration medium; DMSO (the vehicle for the GGG stock solution) was also added to the CXCL12 only condition at a dilution equivalent to the highest GGG concentration used in that experiment. We added 600 μl of these mixtures to a 24-well tissue culture plate. Transwell filters (6 mm insert, 5 μm pore size, Corning) were placed on top of each well, and 100 μL of resensitized cells (2 × 10^5^ cells) was added to the transwell insert. The plate was placed in a 37°C, 5% CO_2_ incubator, and the cells were allowed to migrate for 3 hours, after which the cells in the bottom well were counted by flow cytometry. To determine the degree of migration inhibition, the number of GFP^+^ cells that migrated in each well was normalized to GFP^−^ cells, and this ratio in turn compared to that for CXCL12 alone.

### Proliferation assay with exogenous GGG

P2RY8 KO Ly8 cells were transduced as described in *Individual variant validation* above. Starting on day 3, GGG (or DMSO as a control) was added to the culture media each day to the final desired concentration (ranging from 1 nM to 1 μM GGG, or 1:1000 DMSO). The ratio of GFP^+^ to GFP^−^ cells was tracked over time by flow cytometry at 3, 5, 7, and 10 days after transduction. Cells were passaged at each of those time points as well.

### Candidate ligand migration assay

P2RY8-expressing WEHI-231 cells were produced as previously described^29^. In brief, P2RY8 had been cloned into the murine stem cell virus (MSCV)-GFP retroviral vector. The retrovirus was produced using Plat-E cell line. WEHI-231 cells were transduced in a 6 well plate with retroviral supernatant, centrifuged at 1,340 x g for 2 h at room temperature. The supernatant was removed and standard WEHI-231 culture media replaced. This spinfection was repeated 24 h later.

Post-transduction cells were washed with migration medium (RPMI, 0.5% fatty acid free BSA (Sigma), 10 mM HEPES, and 50 IU penicillin/streptomycin), resuspended at 2 × 10^6^ cells mL^−1^ and resensitized for 10 to 15 minutes at 37°C. Recombinant human CXCL12 (Peprotech) was diluted to 50 ng mL^−1^ in migration medium. GGG or other candidate ligands were diluted to various concentrations in the CXCL12-containing migration medium. We added 600 μl of these mixtures to a 24-well tissue culture plate. Transwell filters (6 mm insert, 5 μm pore size, Corning) were placed on top of each well, and 100 μL of resensitized cells (2 × 10^5^ cells) was added to the transwell insert. The plate was placed in a 37°C, 5% CO_2_ incubator, and the cells were allowed to migrate for 3 hours, after which the cells in the bottom well were counted by flow cytometry. To determine the degree of migration inhibition, the number of GFP+ cells that migrated in each well was normalized to GFP-cells, and this ratio in turn compared to that for CXCL12 alone.

### Phospho-flow

P2RY8 KO Ly8 cells were transduced with variants of interest as described in *Individual variant validation* above. Phospho-flow analyses were performed 5 to 7 days after transduction. Cells were washed in migration medium (RPMI, 0.5% fatty-acid free BSA, 10 mM HEPES, and 50 IU penicillin/streptomycin). They were then resuspended in migration medium at 5 × 10^6^ cells mL^−1^ and resensitized at 37°C for 12 minutes. At that time 100 μL (5 × 10^5^) cells were diluted to 200 uL in 5 mL polystyrene round bottom tube with indicated combinations of 100 nM GGG and 100 ng mL^−1^ CXCL12 and incubated in 37°C water bath for 5 minutes. Afterward, 22 uL of 16% paraformaldehyde was added to each tube and cells fixed at room temperature for 10 minutes, centrifuged, supernatant removed, and 1 mL of cold methanol added while vortexing. The samples were placed at −20°C for 1-2 nights (consistent within given experiment). They were then washed three times with FACS buffer and divided into three equal aliquots, one of which was used for pAkt and one of which was used for pErk. They were blocked for 20 minutes at room temperature with 5% normal goat serum and 1:100 human Fc block (Trustain FcX, Biolegend) and stained at room temperature for 1 hour with a 1:100 dilution of rabbit anti-pAkt (Cell Signaling Technology, Ser473, clone D9E) or 1:100 rabbit anti-pErk1/2 (Cell Signaling Technology, T202/T204, #9102). They were washed twice in FACS buffer, stained for 1 h at room temperature with 1:300 streptavidin-AF647 and 1:250 anti-GFP (Biolegend, clone FM264G), washed with FACS buffer, and analyzed.

### NanoBit β-arrestin assay

This assay was performed in 293T cells. β2AR-LgBiT and β-Arr2-SmBiT plasmids had previously been generated^54^. The β2AR-LgBiT plasmid was digested with EcoRI and BmtI and β2AR replaced with P2RY8 coding sequence, either WT or with V29W, K180I, W181M, or L220Y missense variants, using PCR on the plasmids previously generated for *Individual variant validation*. On day 0, 293T cells were plated into 6-well plate. On Day 1, with cells now ∼80% confluent, cells were transfected. Specifically, to 250 μL Opti-MEM added 400 ng of desired LgBiT construct, 100 ng of β-Arr2-SmBiT, and 1.5 μL of Trans-IT 2020 (Mirus) and gently mixed. After 25 min incubation at room temperature, this was added dropwise to well of the 293T cells. On Day 2, culture medium was removed. Cells were dissociated from plate using PBS with 0.5 mM EDTA, washed, and counted. Cells were resuspended at 4 × 10^5^ cells mL^−1^ in assay solution (Hanks’ balanced salt solution with 20 mM HEPES) and 100 μL (4 × 10^4^ cells) added to wells in 96-well white flat-bottom plate. To this, added 50 μL of 15 μM coelenterazine h (Nanolight). After 10 minutes, measured baseline luminescence. Then added 50 μL of desired GGG dilution in assay solution with 5 μM coelenterazine h to yield final concentrations ranging from 100 pM to 10 μM as well as a DMSO (vehicle) condition. After 10 minutes, measured luminescence again. For analysis, looked at ratio of post-GGG signal to baseline signal, normalized to DMSO condition ratio.

### TRUPATH BRET trimeric G protein assay

This assay was performed in 293T cells. To start, 5 × 10^5^ cells were plated per well in a 6-well plate. The next day, they were transfected. Specifically, to 250 μL of Opti-MEM, added 150 ng of P2RY8 WT, P2RY8 variant, or GFP only plasmid (same POP2E original or modified as used in *Individual variant validation* above), 100 ng pcDNA5/FRT/TO-GAlpha13-RLuc8, 100 ng pcDNA3.1-Beta3, 100 ng of pcDNA3.1-GGamma9-GFP2, and 1.35 μL of Trans-IT 2020 reagent (Mirus) and gently mixed; after 25 minutes incubation at room temperature, this was added dropwise to well of the 293T cells. pcDNA5/FRT/TO-GAlpha13-RLuc8, pcDNA3.1-Beta3, and pcDNA3.1-GGamma9-GFP2 were gifts from Bryan Roth, Addgene plasmid #140986, 140988, and 140991^42^. The next day, 96-well white flat-bottom plates (Greiner Bio-One) were treated for 2 hours with 50 μg mL^−1^ poly-D-lysine (diluted from 1 mg mL^−1^, EMD-Millipore) in sterile ultra-distilled water, washed once with water, then dried for 1 hour. Culture medium removed from the transfected cells, cells were dissociated from plate using PBS with 0.5 mM EDTA, washed, resuspended at 2.67 × 10^5^ cells mL^−1^ in culture medium, and 150 μL (4 × 10^4^ cells) aliquoted per well of poly-D-lysine treated plate. The following day, the plate was centrifuged at 500 x g for 1 min, culture medium removed, and 60 μL of assay medium (Hanks’ balanced salt solution plus 20 mM HEPES) was added to each well. From 2.5 mM stock of coelenterazine 400a (Nanolight), a 50 μM dilution in assay buffer was made. In addition, desired dilutions of GGG in assay were prepared (at four times the ultimately desired concentration). Transported cells on ice to building with BRET-capable plate reader, plate again centrifuged 500 x g for 1 min, and 10 μL of coelenterazine working solution was added. After 10 minutes, 30 μL of desired GGG solution was added, yielding final concentrations ranging from 100 pM to 10 μM; there was also a 1:100 DMSO (vehicle) condition. After 8 minutes, loaded on plate reader and measured 420 ± 20 nm and 515 ± 20 nm for luciferase and GFP, respectively. Measured 6 times, each spaced by 3 minutes. For analysis, calculated luminescence to GFP ratio for each time point and determined mean across the six measurements.

### B cell transduction, transfer, and analysis

P2RY8 was previously cloned into the MSCV-GFP retroviral vector as described^29^. The GFP was replaced by digestion of MSCV-P2RY8-IRES-GFP with BstXI and PacI and insertion of TagBFP PCR product using HiFi DNA Assembly Kit (NEB). The WT P2RY8 MSCV was digested with BglII and NotI and K180I or W181M P2RY8 PCR products (from the constructs detailed in *Individual variant validation* above) were inserted using HiFi DNA Assembly Kit (NEB). Retrovirus encoding P2RY8-TagBFP, K180I-GFP, and W181M-GFP were produced using Plat-E packaging line as described^28^. EasySep kits were used to enrich B cells from mouse spleens by removing T cells with biotin-conjugated anti-CD3ε (Biolegend, clone 145-2C11) and streptavidin-conjugated beads (EasySep Streptavidin RapidSpheres). B cells were cultured in complete RPMI (10% FBS, 10 mM HEPES, 1x GlutaMax, 50 IU penicillin/streptomycin, and 55 μM β-mercaptoethanol) along with 25 μg mL^−1^ (1:4000 dilution) anti-CD180 (BD Biosciences, clone RP/14).

Transduction was performed 24 hours after activation. Plates were centrifuged and culture supernatant was saved. Retroviral supernatants were thawed from −80°C and HEPES added to 20 mM and polybrene (EMD Millipore) added to 2 μg mL^−1^. This was used to spinfect the cells for 2 hours at room temperature, centrifuging at 2500 rpm. The viral supernatant was then aspirated and half the original culture medium was replaced along with an equal volume of culture medium with freshly added anti-CD180. The spinfection was repeated a second time 24 hours later.

Twenty-four hours after the second transduction, cells were collected from the plate, washed twice, and placed on ice. A small aliquot was analyzed by flow cytometry to determine the proportion of transduced (GFP^+^ or TagBFP^+^) cells. Cells were diluted and pooled (WT P2RY8 and K180I P2RY8 or WT P2RY8 and W181M P2RY8) to desired ratio and injected IV into recipient mice, who had been immunized intraperitoneally 7 days earlier with 2 × 10^8^ SRBCs. Mice received 4-5 × 10^6^ WT-TagBFP^+^ cells and 7-10 × 10^6^ K180I-GFP^+^ or W181M-GFP^+^ cells.

Mice were analyzed 24 hours after transfer. Spleens were harvested, cut into 4-5 sections, and fixed for 30 minutes in 4% paraformaldehyde in PBS at 4°C, washed, then dehydrated overnight at 4°C in 30% sucrose solution. Spleen segments were placed in cassette with OCT medium (Sakura), flash frozen, and placed at −80°C for storage. Cryostat was used to cut 7 μm sections. These were dried at room temperature for one hour then rehydrated with PBS 0.1% fatty-acid-free (FAF) BSA for 10 minutes. They were stained overnight at 4°C with primary stain of PBS 0.1% FAF-BSA, 1:100 normal mouse serum, 1:100 anti-GFP AF488 (rabbit polyclonal, Invitrogen A21311), 1:100 anti-CR1 (CD35)-biotin (BD 553816, clone 8C12), 1:100 anti-IgD AF647 (Biolegend 405708, clone 11-26c2a), and 1:200 or 1:250 anti-TagFP AZ568 (NanoTag N0502-AF568). Cells were washed 3 times with PBS 0.1% FAF-BSA then secondary stain was PBS 0.1% FAF-BSA, 1:100 normal mouse serum, 1:200 streptavidin-AMCA for 1 hour at room temperature. Cells were washed 2 times with PBS 0.1% FAF-BSA, once with PBS, then cover slip affixed using Fluoro-Mount, allowed to harden at 4 C for one hour. Images were acquired using Zeiss AxioObserver Z1 inverted microscope.

Initial image processing performed in ZEN (Zeiss): AMCA channel black/white rescaled from 0-16,034 to 1,596-16,034 for all images obtained. AF555 channel black/white rescaled from 0-16,034 to 1,047-9,052 all images in figures. Further analysis performed using Fiji^55^. Follicles were cropped from larger images and regions of interest containing GCs defined. Using consistent thresholds across images, GFP and BFP signal was made binary and Analyze Particles function (size 10-150 μm^2^, circularity 0.3-1) used to count cells within follicles and within subsections of follicles coinciding with GCs. Analysis included at least 45 GCs from each mouse.

### Bone marrow chimera generation and analysis

Mice to be used as bone marrow (BM) donors (C57BL/6J, 8-12 weeks old) were injected intraperitoneally with 3 mg 5-fluorouracil (Sigma). BM was collected after 4 days and cultured in DMEM containing 15% FBS, penicillin (50 IU mL^−1^), streptomycin (50 μg mL^−1^), and 10 mM HEPES, supplemented with 250 ng mL^−1^ IL-3, 50 ng mL^−1^ IL-6, and 100 ng mL^−1^ SCF (Peprotech). Cells were transduced with MSCV-EV-GFP, MSCV-P2RY8(WT)-GFP, MSCV-P2RY8(K180I)-GFP, or MSCV-P2RY8(W181M)-GFP) on days 1 and 2 using viral supernatant concentrate thawed, diluted to original concentration with DMEM, 10% FBS, 10 mM HEPES, and 4 ug mL^−1^ polybrene and transduction performed via 2 hr centrifugation at 2400 rpm at 32°C, followed by replacement with culture media. Recipient CD45.1 B6 mice, age 7-10 weeks, were lethally irradiated with 900 rads (evenly split dose separated by 3 h) and then injected IV with relevant transduced BM cells. For every 3-3.5 recipient mice, 1 donor mouse was used. Mice were analyzed 10-12 weeks after reconstitution.

For analysis, spleen, mesenteric lymph nodes, and Peyer’s patches were isolated. Mesenteric lymph nodes and Peyer’s patches were digested for 20 min at 37°C, shaking, in 1 mg mL^−1^ collagenase VIII (Sigma) in RPMI with 10% FBS. After mashing, straining, and washing, tissues were blocked for 10 min with 1:100 Fc block (Bio-X-Cell) in FACS buffer (PBS, 2% FBS, 1 mM EDTA), then stained for 30 minutes on ice with anti-CD45.2 PE (Biolegend 109808), anti-CD45.1 APC (Cytek 20-0453-U100), anti-GL7 PacBlue (Biolegend 144614), anti-IgD PerCP-Cy5.5 (Biolegend 405710), anti-CD95 PE-Cy7 (Fisher 557653), anti-B220 BV785 (Biolegend 103246), and, for spleen, also anti-CD8a BV510 (Biolegend 100752), anti-CD11b BV570 (Biolegend 101233), anti-CD4 BV711 (Biolegend 100447), and anti-CD3ε AF700 (Biolegend 152316), all 1:200. Cells were washed, stained for 10 minutes with e780 fixable viability dye (eBiosciences), and washed again. Cells were then analyzed on FACSymphony.

### Expression and purification of P2RY8-miniG_13_ protein

Human P2RY8 (Uniprot: Q86VZ1) was cloned into pCDNA-Zeo-TetO, a custom pcDNA3.1 vector containing a tetracycline-inducible gene expression cassette^56^. The resulting P2RY8 construct comprised an N-terminal influenza haemagglutinin signal sequence followed by a M1-FLAG (DYKDDDD) epitope tag. The P2RY8 construct was furthermore fused at the C-terminus to the miniGα_13_ protein via a linker sequence containing a human rhinovirus 3C (HRV 3C) protease cleavage site. The resulting P2RY8–miniGα_13_ construct was transiently transfected into inducible Expi293F TetR cells (unauthenticated and untested for mycoplasma contamination; Thermo Fisher) using the ExpiFectamine 293 Transfection Kit (Thermo Fisher) following the manufacturer’s instructions. After 16 h, protein expression was induced with 1 µg ml^−1^ doxycycline hyclate (Sigma Aldrich), and the culture was placed in a shaking incubator maintained at 37°C and 8% CO_2_. After 36 h, cells were harvested by centrifugation at 4,000 x g and stored at −80°C.

For receptor purification, cells were thawed and resuspended in lysis buffer comprising 20 mM HEPES pH 7.50, 10 µM S-geranylgeranyl-L-glutathione (GGG, Cayman Chemical), 100 µM tris(2-carboxyethyl)phosphine (TCEP; Fischer Scientific), and a Pierce protease inhibitor tablet (Thermo Scientific). Cells were lysed by stirring for 15 min at 4 °C and harvested by centrifugation at 16,000 x g for 15 min. To increase the fraction of GGG-bound P2RY8, we also reconstituted GGG into detergent to overcome its low solubility. 25 mg of GGG and 1.5 g of lauryl maltose neopentyl glycol (L-MNG; Anatrace) was dissolved in methanol and allowed to mix for 10 min at RT. The LMNG-GGG mixture was was placed under an Argon stream evaporated to dryness followed by vaccuum dessication overnight. The dry LMNG-GGG mixture was resuspended in 20 mM HEPES pH 7.50, 100 mM NaCl by sonication. Cell pellet was dounce-homogenized in ice-cold solubilization buffer comprising 50 mM HEPES, pH 7.5, 150 mM NaCl, 1% (w/v) lauryl maltose neopentyl glycol (L-MNG; Anatrace) containing 3 mol% GGG, 0.1% (w/v) cholesteryl hemisuccinate (CHS, Steraloids), 5 mM adenosine 5′-triphosphate (ATP; Fischer Scientific), 2 mM MgCl_2_, 100 µM TCEP, 10 µM GGG, and a Pierce protease inhibitor tablet (Thermo Scientific). The sample was stirred for 1 h at 4°C, and the detergent-solubilized fraction was clarified by centrifugation at 20,000 x g for 30 min. The detergent-solubilized sample was supplemented with 4 mM CaCl_2_ and incubated in batch with homemade M1-FLAG-antibody conjugated CNBr-Sepharose under slow rotation for 1 h at 4 °C. The P2RY8-bound resin was transferred to a glass column and washed with 20 ml of ice-cold buffer comprising 50 mM HEPES, pH 7.5, 150 mM NaCl, 0.1% (w/v) L-MNG containing 3 mol% GGG, 0.01% (w/v) CHS, 5 mM ATP, 2 mM CaCl_2_, 2 mM MgCl_2_, 100 µM TCEP. This was followed by 10 column volumes of ice-cold 50 mM HEPES, pH 7.5, 150 mM NaCl, 0.0075% (w/v) L-MNG containing 3 mol% GGG, 0.0025% (w/v) glyco-diosgenin (GDN; Anatrace), 0.001% (w/v) CHS, 2 mM CaCl2, 100 µM TCEP, and 10 µM GGG. Receptor-containing fractions were eluted with ice-cold 20 mM HEPES, pH 7.5, 150 mM NaCl, 0.0075% (w/v) L-MNG containing 3 mol% GGG, 0.0025% (w/v) GDN, 0.001% (w/v) CHS, 5 mM EDTA, 0.2 mg ml^−1^ FLAG peptide, 100 µM TCEP, and 10 µM GGG. Fractions containing P2RY8– miniGα_13_ were concentrated in a 50-kDa MWCO spin filter (Amicon) and purified over a Superdex 200 Increase 10/300 GL (Cytiva) size-exclusion chromatography (SEC) column, which was equilibrated with 20 mM HEPES, pH 7.5, 150 mM NaCl, 0.0075% (w/v) L-MNG containing 3 mol% GGG, 0.0025% (w/v) GDN, 0.001% (w/v) CHS, 100 µM TCEP, and 10 µM GGG. Fractions containing monodisperse P2RY8–miniGα_13_ were combined and concentrated in a 50-kDa MWCO spin filter before complexing with Gβ_1_γ_2_.

### Expression and purification of Gβ_1_γ_2_

Purified Gβ_1_γ_2_ was generated as described previously^57^. Briefly, a baculovirus was generated in Spodoptera frugiperda Sf9 insect cells (unauthenticated and untested for mycoplasma contamination, Expression Systems) using the pVLDual expression vector encoding both human Gβ _1_ subunit with a HRV 3C cleavable N-terminal 6xHis-tag and the untagged human Gγ_2_ subunit. Gβ_1_γ_2_ was expressed in Trichoplusia ni Hi5 insect cells (unauthenticated and untested for mycoplasma contamination, Expression Systems) by infection with Gβ_1_γ_2_-baculovirus at a density of 3.0 × 10^6^ cells ml^−1^ and grown for 48 h at 27°C with 130 r.p.m. shaking. Harvested cells were resuspended in lysis buffer comprised of 20 mM HEPES, pH 8.0, 5 mM β-mercaptoethanol (β-ME), 20 µg ml^−1^ leupeptin, and 160 µg ml^−1^ benzamidine. Lysed cells were pelleted at 20,000 x g for 15 min, and solubilized with 20 mM HEPES, pH 8, 100 mM sodium chloride, 1% (w/v) sodium cholate (Sigma Aldrich), 0.05% (w/v) n-dodecyl-β-d-maltopyranoside (DM; Anatrace) and 5 mM β-ME. Detergent-solubilized Gβ_1_γ_2_ was clarified by centrifugation at 20,000 x g for 30 min and was then incubated with HisPur Ni-NTA resin (Thermo Scientific) under slow rotation for 1.5 h at 4°C. Gβ_1_γ_2_-bound resin was transferred to a glass column and washed extensively. Detergent was exchanged on-column to 0.1% (w/v) L-MNG and 0.01% (w/v) CHS, and Gβ_1_γ_2_ was eluted with 20 mM HEPES pH 7.50, 100 mM NaCl, 0.1% (w/v) L-MNG, 0.01% (w/v) CHS, 300 mM imidazole, 1 mM DL-dithiothreitol (DTT), 20 µg ml^−1^ leupeptin and 160 µg ml^−1^ benzamidine. Gβ_1_γ_2_-containing fractions were pooled and supplemented with homemade 3C protease before overnight dialysis in buffer comprised of 20 mM HEPES, pH 7.50, 100 mM NaCl, 0.02% (w/v) L-MNG, 0.002% (w/v) CHS, 1 mM DTT and 10 mM imidazole. Cleaved Gβ_1_γ_2_ was isolated by reverse Ni-NTA, and dephosphorylated by treatment with lambda phosphatase (New England Biolabs), calf intestinal phosphatase (New England Biolabs) and antarctic phosphatase (New England Biolabs) for 1 h at 4 °C. The geranylgeranylated Gβ_1_γ_2_ heterodimer was isolated by anion exchange chromatography using a MonoQ 4.6/100 PE (Cytiva) column, before overnight dialysis in 20 mM HEPES, pH 7.5, 100 mM NaCl, 0.02% (w/v) L-MNG and 100 µM TCEP. The final sample was concentrated on a 3-kDa MWCO spin filter (Amicon), and 20% (v/v) glycerol was added before flash freezing in liquid N2 for storage at −80°C.

### Preparation of the active-state of P2RY8-G_13_ complex

To prepare the P2RY8–G_13_ complex, a twofold molar excess of purified Gβ_1_γ_2_ was added to SEC-purified P2RY8–miniGα_13_ followed by overnight incubation on ice. The sample was purified on a Superdex 200 Increase 10/300 GL SEC column, equilibrated with 20 mM HEPES, pH 7.5, 150 mM NaCl, 0.0075% (w/v) L-MNG containing 3 mol% GGG, 0.0025% (w/v) GDN, 0.001% (w/v) CHS and, 30 µM GGG. Fractions containing the monomeric P2RY8–G13 heterotrimeric complex were concentrated on a 50-kDa MWCO spin filter (Amicon) immediately before cryo-EM grid preparation.

### Cryo-EM vitrification, data collection, and processing

2.75 µl of the purified P2RY8–G13 complex was applied at 1.4 mg ml^−1^ to glow-discharged 300 mesh R1.2/1.3 UltrAuFoil Holey gold grids (Quantifoil). Grids were plunge-frozen in liquid ethane using a Vitrobot Mark IV (Thermo Fisher) with a 10-s hold period, blot force of 0, and blotting time varying between 1.5 and 3.0 s while maintaining 100% humidity and 4°C. Vitrified grids were clipped with Autogrid sample carrier assemblies (Thermo Fisher) immediately before imaging. Movies of P2RY8–G_13_ embedded in ice were recorded on a 300 kV FEI Titan Krios microscope equipped with a Falcon 4i camera (Thermo Scientific, located at the HHMI Janelia Research Campus). A nominal magnification of × 130,000 was used in resolution mode with a physical pixel size of 0.94 Å per pixel, and movies were recorded with a total exposure of 50 e− Å^−2^. Movies (n = 12,053) were imported into cryoSPARC (Structura Biotechnology) with 80 EER fractions, motion-corrected micrographs were generated with the patch motion correct tool beofre calculation of patch contrast transfer functions (patch CTFs). A threshold of CTF fit resolution of more than 6 Å was used to exclude low-quality micrographs. Particles were template picked using a 20 Å low-pass-filtered model that was generated ab initio from data collected during earlier screening sessions on a 200-kV Glacios microscope located at UCSF. Particles (n = 10,117,438) were extracted with a box size of 288 pixels binned to 72 pixels and sorted by two rounds of 2D classification. The resulting 2,352,975 particles were sorted by ab initio 3D reconstruction with two classes, re-extracted with a box size of 288 pixels binned to 144 pixels, and sorted by additional two rounds of ab initio 3D construction. Particles (n = 799,516) were extracted with an unbinned box size of 288 pixels and were subjected to non-uniform refinement followed by local refinement using a mask covering only the 7TM domain of P2RY8. Particles were further sorted by two rounds of 3D classification using 4 classes and a filter resolution of 3.00 Å and 2.75 Å respectively. The resulting 90,243 particles were subjected to non-uniform refinement followed by local refinement using an inclusion mask covering the 7TM domain, using poses/shift Gaussian priors with standard deviation of rotational and shift magnitudes limited to 1° and 1 Å, respectively.

### Model building and refinement

Model building and refinement were carried out using a starting model based on the AlphaFold2 predicted structure of P2RY8 (Uniprot: Q86VZ1), the deposited structure of Gα_13_ (PDB code: 7T6B)^58^, and the deposited structure of Gβ_1_γ_2_ (PDB code: 8F76)^57^ which were fitted into the P2RY8–G_13_ map using UCSF ChimeraX. A draft model was generated using ISOLDE^59^ and was further refined by iterations of real-space refinement in Phenix v1.20^60^ and manual refinement in Coot v0.8.9.2^61^. The GGG model and rotamer library were generated with the eLBOW extension^62^ in Phenix and docked using Coot. The resulting model was extensively refined in Phenix and map-model validations were carried out using Molprobity v4.5 and EMRinger^63^.

### Chemical synthesis

GGG used in WEHI migration assay was synthesized in house as previously described^29^. GGG for all other assays was purchased from Cayman Chemical and resuspended in DMSO at 1 mM to form stock solution. Leukotriene C_4_ was also purchased from Cayman Chemical.

Synthesis of other glutathione conjugates followed similar approach to GGG^29^. Compound 1 refers to S-octane-L-glutathione, compound 2 refers to S-hexadecane-L-glutathione, compound 3 refers to S-isoprenyl-L-glutathione, and compound 4 refers to S-farnesyl-L-glutathione. Unless otherwise stated, all reagents were purchased from MilliporeSigma. Briefly, 20 mg (1 eq.) of farnesol (for compound 4) was stirred in 1 ml of dry DCM under an atmosphere of nitrogen at room temperature. Triphenylphospine (1.3 eq.) was added, followed by carbon tetrabromide (1.3 eq.), and the reaction was stirred for a further 2–4 hours at room temperature. After concentrating the crude reaction under reduced pressure, a small volume of n-hexane was added and the resulting precipitate removed by filtration. Concentration, precipitation and filtration was repeated once again and the concentrated filtrate used in the next step without further purification.

Glutathione (1.1 eq) was dissolved in the minimal volume of 2 M NaOH, and ethanol added until the solution started to become cloudy. Either the compound generated in the first step (for compound 4), or 1-bromooctane, 1-bromohexadecane, or 3,3-Dimethylallyl bromide (for compounds 1, 2, and 3) (1 eq.) was added dropwise and the reaction stirred at room temperature overnight. The pH of the reaction mixture was reduced to 2 by addition of 1 M HCl, and the reaction cooled in an ice bath until a precipitate formed. This precipitate was collected by filtration, washed with a small volume of ice-cold ethanol and then ice-cold water, and dried to yield the final glutathione conjugate.

### Statistics and graphing

Bar and line graphs and included statistics made using Prism (GraphPad). Other plots and statistics using R version 4.3.0 in RStudio 2023.03.0+386, relevant packages tidyverse 2.0.0, ggplot2 3.4.2. P2RY8 structure analysis using ChimeraX version 1.8^64,65^. Regression analysis using Jupyter Notebook 6.5.4, Python 3.11.4, numpy 1.24.3, pandas 1.5.3, scipy 1.10.1, and sklearn 1.5.2.

## Data Availability

Sequencing data from the deep mutational scan have been deposited in the NCBI Sequence Read Archive under BioProject PRJNA1179413 and are publicly available as of the date of publication. Coordinates for the P2RY8-Gα_13_ complex have been deposited in the RCSB Protein Data Bank under accession codes 9ECJ. EM density map for the P2RY8-Gα_13_ complex has been deposited in the Electron Microscopy Data Bank under accession codes EMD-47912 (full map) and EMD-47914 (7TM map). *Previously available data:* Germline variants from gnomAD^2^ v4.0.0 at https://gnomad.broadinstitute.org. Non-hematologic cancer variants and some DLBCL / Burkitt lymphoma variants from COSMIC^45^ at https://cancer.sanger.ac.uk/cosmic, accessed Feb. 1, 2024.

## Code Availability

Analysis code are available on GitHub at https://github.com/yelabucsf/P2RY8_DMS. Any additional information required to reanalyze the data reported in this paper is available from the lead contact upon request.

## Acknowledgments

T.N.L. is a fellow in the Pediatric Scientist Development Program (NICHD K12-HD000850), and also supported by the UCSF Center for Rheumatic Diseases. J.G.C. is an investigator of the Howard Hughes Medical Institute. This work was supported in part by NIH grant R01 AI045073. C.J.Y. is supported by the NIH grants R01AI171184, P01AI172523, and the Arc Research Institute, and is a member of the Gladstone-UCSF Institute of Genomic Immunology (GIGI) and Parker Institute for Cancer Immunotherapy (PICI). We acknowledge the PFCC (RRID:*SCR_018206*) supported in part by Grant NIH P30 DK063720 and by the NIH S10 Instrumentation Grant S10 1S10OD021822-01. Sequencing was performed at the UCSF CAT, supported by UCSF PBBR, RRP IMIA, and NIH 1S10OD028511-01 grants. Portions of this work were performed on the Wynton HPC Co-Op cluster, which is supported by UCSF research faculty and UCSF institutional funds. The authors wish to thank the UCSF Wynton team for their ongoing technical support of the Wynton environment. We thank Drs. Li Wang, Glen Gilbert, and David Bulkley at the UCSF Bay Area cryo-EM consortium for help in Glacios microscope operation. Cryo-EM equipment at UCSF is partially supported by NIH grants S10OD020054, S10OD021741 and S10OD026881 and Howard Hughes Medical Institute. We thank Drs. Shinxin Yang and Rui Yan at the HHMI Janelia CryoEM Facility for help in Krios microscope operation and data collection. Molecular graphics and analyses performed with UCSF ChimeraX, developed by the Resource for Biocomputing, Visualization, and Informatics at the University of California, San Francisco, with support from National Institutes of Health R01-GM129325 and the Office of Cyber Infrastructure and Computational Biology, National Institute of Allergy and Infectious Diseases. We thank Zachary Earley for assistance with murine tissue analysis.

## Author contributions

T.N.L., C.B.B., A.M., J.C.G., and C.J.Y. conceived and designed this study. T.N.L. performed DMS and follow-up experiments, performed most VEP analyses, and took lead in writing the paper. C.B.B. performed cryo-EM sample preparation and data analysis. J.C.G., C.J.Y, A.M., and C.B.B. co-wrote the paper. T.D. and V.N.. performed ESM1b fine-tuning and some related analysis. F.D.W. synthesized candidate ligands. E.L. performed migration assay with candidate ligands. J.A. participated in creating the mouse chimeras. Y.X. participated in cloning and plasmid preparation. A.S. participated in NanoBit and TRUPATH assays and cloning and plasmid preparation. T.M. and N.B. assisted with VEP analysis.

## Ethics Declaration

A.M. is a founder of Epiodyne and Stipple Bio, consults for Abalone, and serves on the scientific advisory board of Septerna and Alkermes. J.G.C. is a Scientific Advisory Board member of Be Biopharma and consults for Lycia Therapeutics and DrenBio Inc. C.J.Y. is founder for and holds equity in DropPrint Genomics (now ImmunAI) and Survey Genomics, a Scientific Advisory Board member for and hold equity in Related Sciences and ImmunAI, a consultant for and hold equity in Maze Therapeutics, and a consultant for TReX Bio, HiBio, ImYoo, and Santa Ana. Additionally, C.J.Y is also newly an Innovation Investigator for the Arc Institute. C.J.Y. has received research support from Chan Zuckerberg Initiative, Chan Zuckerberg Biohub, Genentech, BioLegend, ScaleBio, and Illumina.

