## Extended Data Figures for "Phenotypic pleiotropy of missense variants in human B cell-confinement receptor P2RY8"

### Extended Data Figure 1

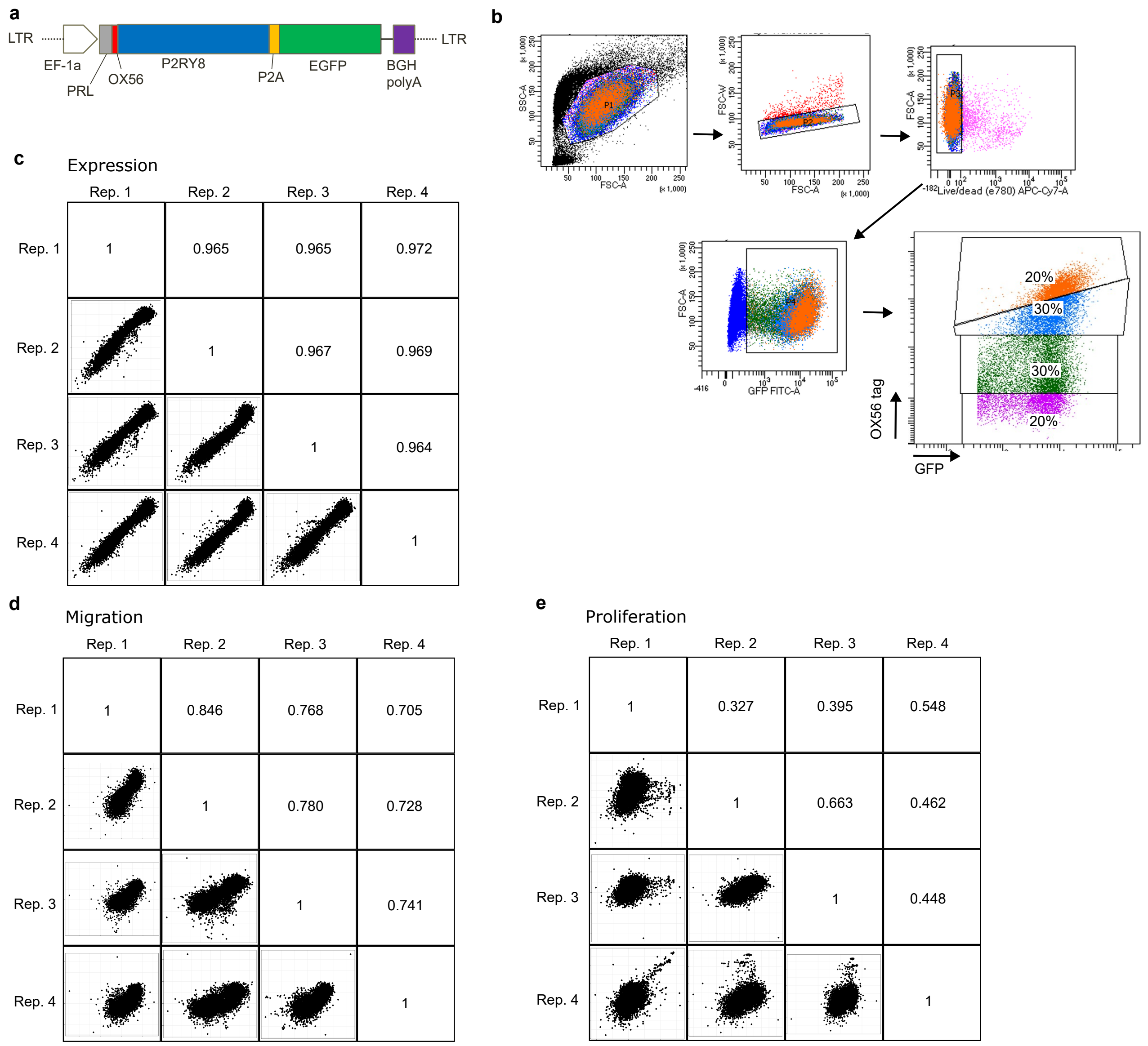

**Extended Data Fig. 1: DMS approach. a,** Schematic of lentiviral vector used for deep mutational scanning approach, OX56 is a peptide tag[66], PRL is preprolactin signal peptide. **b,** Representative flow-assisted cell sorting gating used for surface expression portion of screen. **c, d, e,** Dot plots and Pearson correlations of Enrich2 variant scores for each combination of the four replicates for **(c)** surface expression, **(d)** migration, and **(e)** proliferation.

Extended Data Figure 2

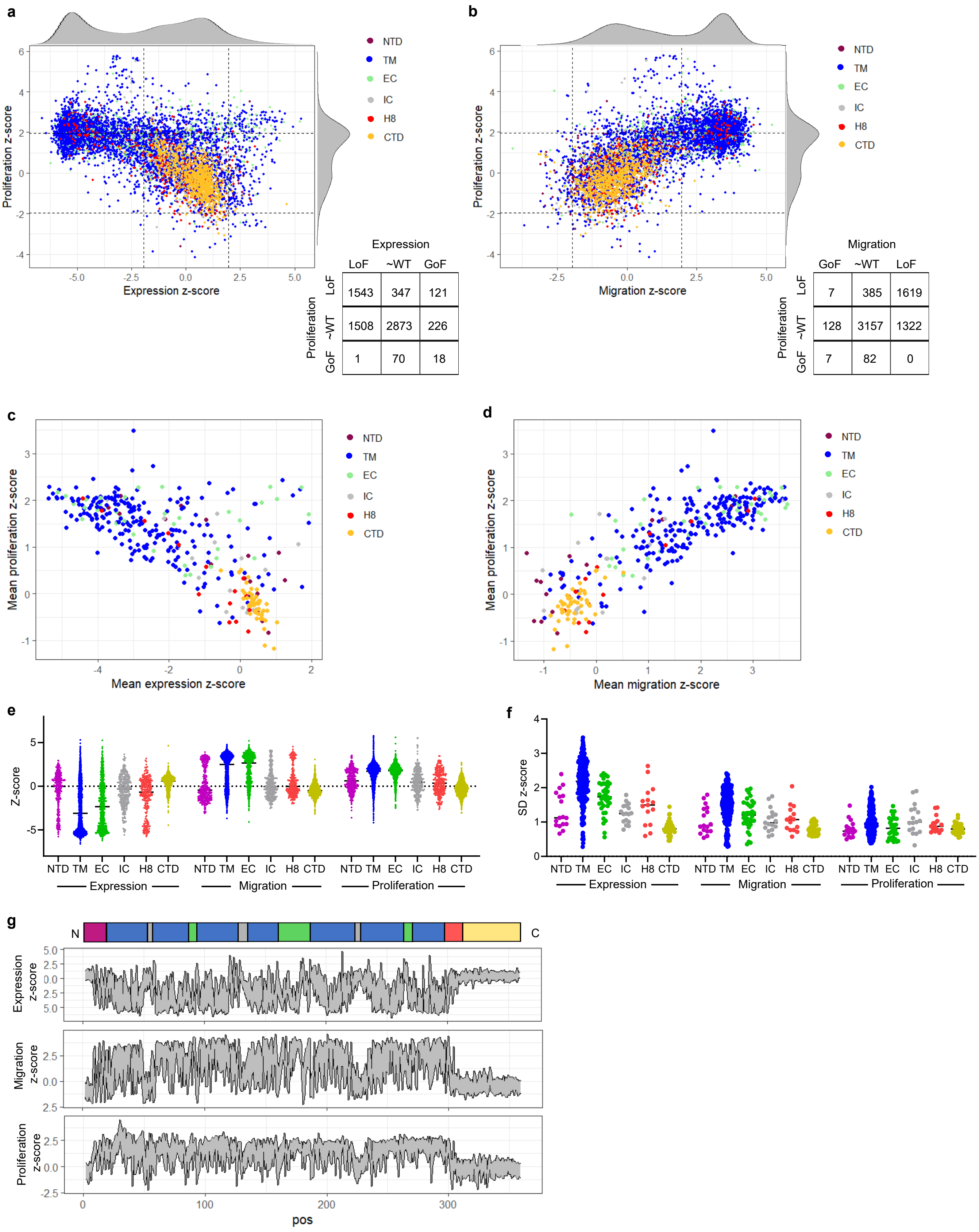

**Extended Data Fig. 2: Heterogeneity of P2RY8 DMS results. a,** Plot comparing expression and proliferation z-scores of missense variants, colored by protein domain, with table showing variant count by section, boundaries at z-scores of 2 and -2. **b,** Plot comparing migration and proliferation z-scores of missense variants, colored by protein domain, with table showing variant count by section, boundaries at z-scores of 2 and -2. **c,** Plot comparing expression and proliferation mean z-scores for each position, colored by protein domain. **d,** Plot comparing migration and proliferation mean z-scores for each position, colored by protein domain. **e,** Variant expression, migration, and proliferation z-scores partitioned by protein domain; lines mark medians; negative z-scores are LoF for surface expression, GoF for migration and proliferation. **f,** Standard deviation of variant expression, migration, and proliferation z-scores within each position, partitioned by protein domain; lines mark medians. **g,** Line plots depicting maximum and minimum variant z-scores at each position for expression, migration, and proliferation. CTD, C-terminal domain; EC, extracellular loop; GoF, gain-of-function; H8, helix 8; IC, intracellular loop; LoF, loss-of-function;  N, N-terminus; NTD, N-terminal domain; TM, transmembrane helices.

### Extended Data Figure 3

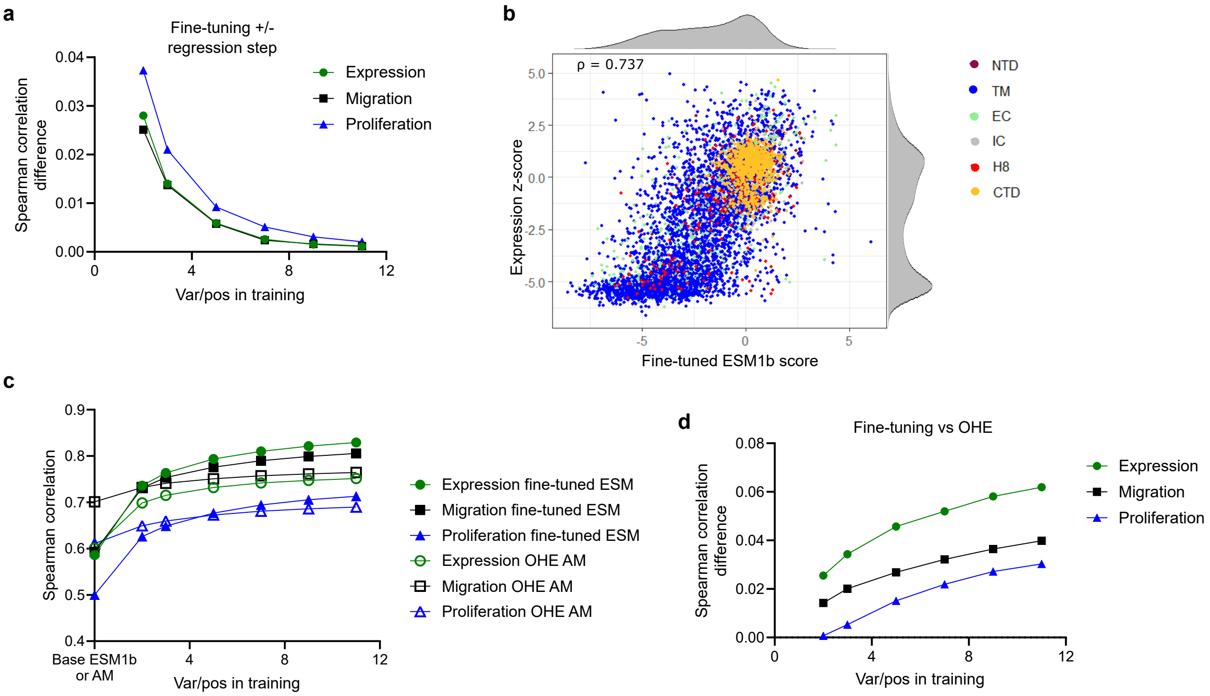

**Extended Data Fig. 3: Supplemental ESM1b fine-tuning analyses. a,** Plot comparing mean Spearman correlations between DMS effect sizes and fine-tuned ESM1b with various training set sizes, with or without final ridge regression step (value > 0 indicates higher correlation with ridge regression). **b,** Plot comparing representative 2 variant/position fine-tuned ESM1b and DMS expression z-score, variants colored by protein domain. **c,** Plot comparing Spearman correlation between DMS z-scores and fine-tuned ESM1b or OHE AM scores as training set size varies. Shows mean and SD (k = 50). **d,** Plot comparing difference in mean Spearman correlation at given training set size between DMS z-scores and fine-tuned ESM1b or. OHE ESM1b (values > 0 indicate higher correlation with fine-tuning). CTD, C-terminal domain; EC, extracellular loop; GoF, gain-of-function; H8, helix 8; IC, intracellular loop; LoF, loss-of-function; NTD, N-terminal domain; OHE, one-hot encoded regression; TM, transmembrane helices; ρ, Spearman correlation coefficient.

### Extended Data Figure 4

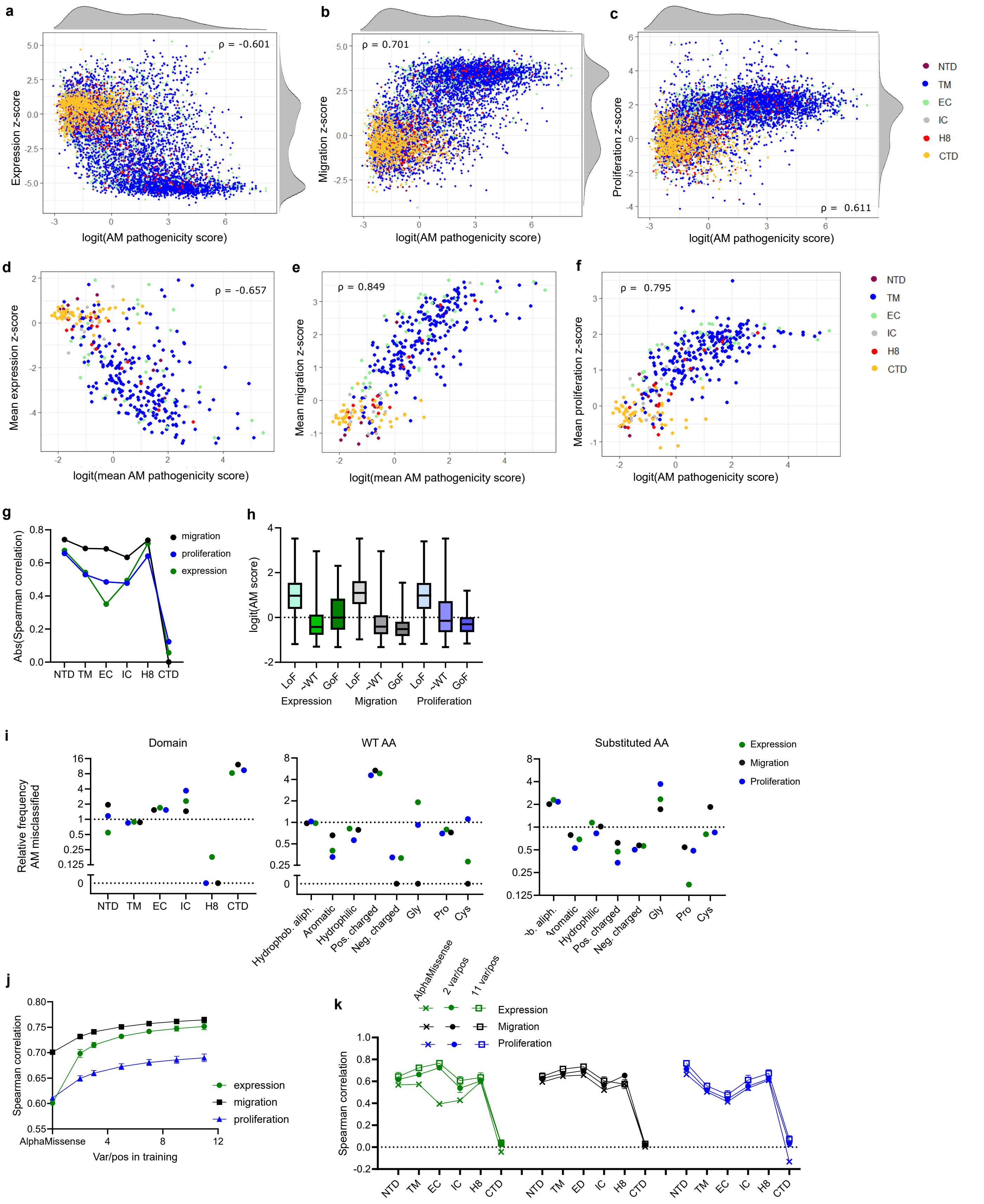

**Extended Data Fig. 4: Improving AlphaMissense variant effect prediction using limited experimental data. a-c** Plots comparing AM pathogenicity score and **(a)** expression z-score, **(b)** migration z-score, or **(c)** proliferation z-score for each variant, colored by protein domain. **d-f,** Plots comparing mean AM pathogenicity score and **(d)** expression mean z-score, **(e)** migration mean z-score, or **(f)** proliferation mean z-score for each position, colored by protein domain. **g,** Plot of Spearman correlation between logit-transformed AM pathogenicity score and variant expression, migration, and proliferation z-scores, variants partitioned by protein domain. **h,** Plot comparing distribution of AM pathogenicity scores for each variant, partitioned by DMS phenotypes. Whiskers extend to maximum and minimum, boxes shows 25th percentile, median, and 75th percentile. **i,** Plots showing proportion of deleterious variants (z-score < -2 for expression, > 2 for migration and proliferation) with benign AM pathogenicity scores (< 0.34) relative to proportion of deleterious variants with non-benign AM scores, as partitioned by protein domain (left), WT amino acid class (center), and substituted (variant) amino acid class (right). Hydrophobic aliphatic residues are A, I, L, M, and V; aromatic residues are F, W, and Y; hydrophilic residues are N, Q, S, and T; positively charged residues are H, K, and R; negatively charged residues are D and E. **j,** Plot of Spearman correlation between OHE AM scores and DMS z-scores as training set size varies. Shows mean and SD (k = 50). **k,** Plot comparing Spearman correlation when using AM score or OHE AM scores with 2 variants/position or 11 variants/position training sets, partitioned by domain. Shows means and SD (k = 50). AM, AlphaMissense; CTD, C-terminal domain; EC, extracellular loop; GoF, GoF, gain-of-function, z-score > 2 for expression, < -2 for migration, proliferation; LoF, loss-of-function, z-score < -2 for expression, > 2 for migration, proliferation; H8, helix 8; IC, intracellular loop; NTD, N-terminal domain; OHE, one-hot encoded regression, ρ, Spearman correlation coefficient.

Extended Data Figure 5

**a**

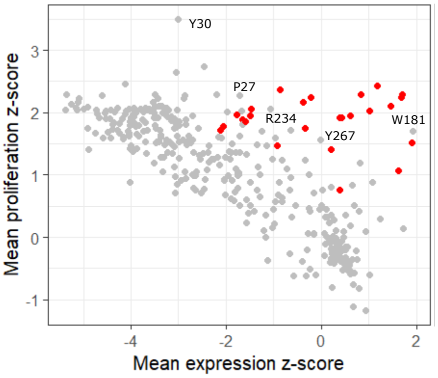

**Extended Data Fig. 5: Proliferation phenotype of high expression-adjusted migration score positions.** **a,** Plot comparing expression and proliferation mean z-scores for each position, with those with high expression-adjusted migration scores (Fig. 3a) colored red; select additional variants labeled.

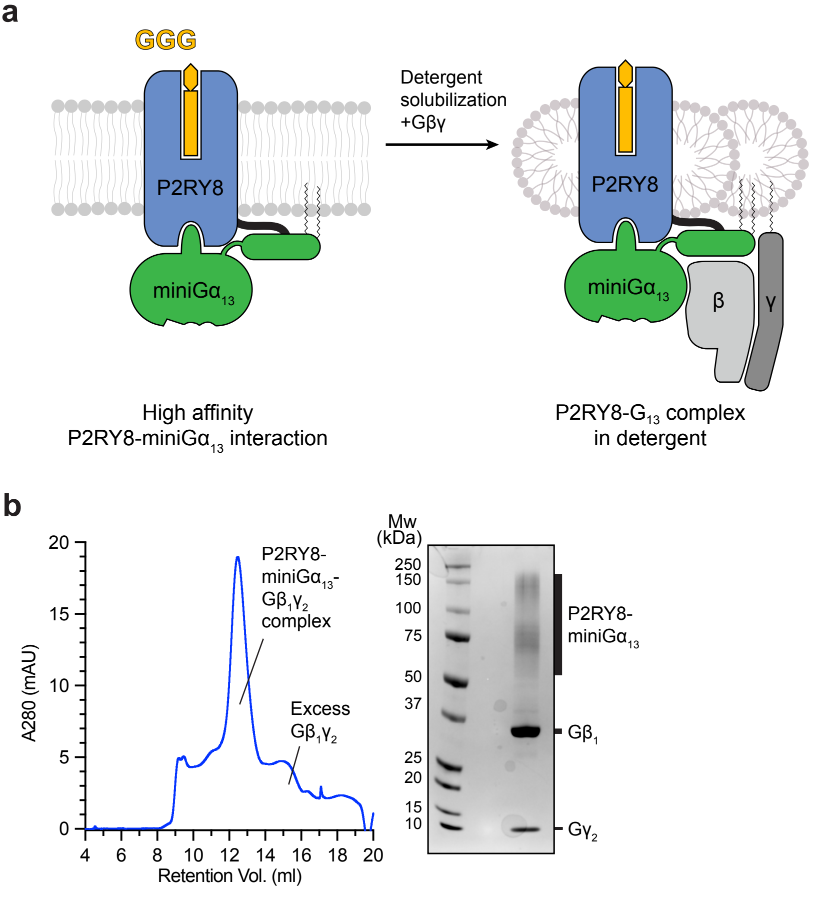

a
GGG
P2RY8
miniGα13
Detergent
solubilization
+Gβγ
P2RY8
miniGα13
β
γ
High affinity
P2RY8-miniGα13 interaction
P2RY8-G13 complex
in detergent
b
A280 (mAU)
Retention Vol. (ml)
P2RY8-
miniGα13-
Gβ1γ2
complex
Excess
Gβ1γ2
Mw
(kDa)
250
150
100
75
50
37
25
20
15
10
P2RY8-
miniGα13
Gβ1
Gγ2

**Extended Data Fig. 6: Biochemical preparation of P2RY8-$G_{13}$ complex bound to GGG. a,** Schematic outlining strategy for stabilization and purification of P2RY8 bound to $G_{13}$ and GGG. **b,** Size-exclusion chromatogram of purified P2RY8-$G_{13}$ complex used for structure determination together with representative SDS-PAGE gel analysis of the collected fraction containing the P2RY8-$G_{13}$ complex.

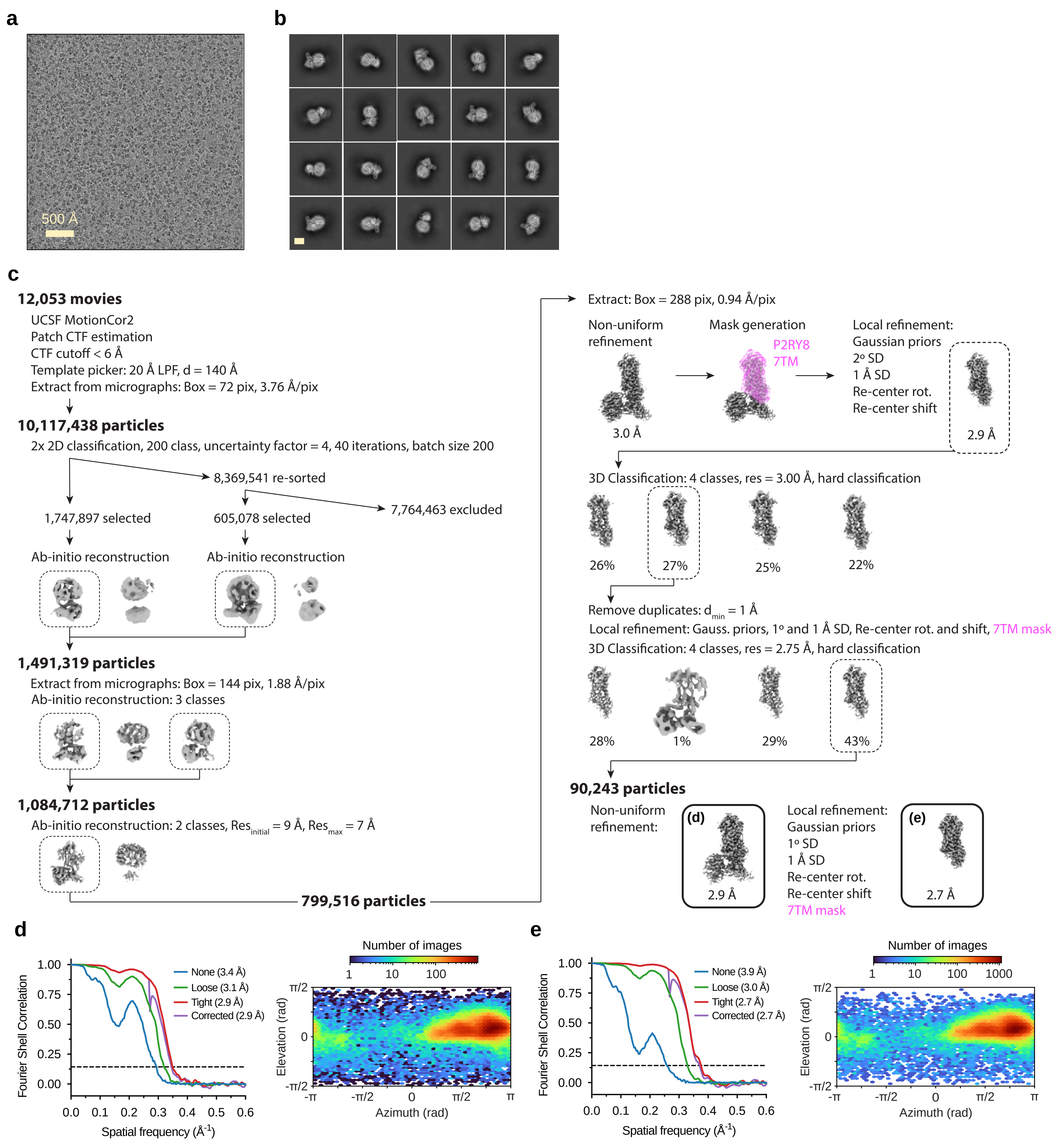
a
500 Å
b
c
12,053 movies
UCSF MotionCor2
Patch CTF estimation
CTF cutoff < 6 Å
Template picker: 20 Å LPF, d = 140 Å
Extract from micrographs: Box = 72 pix, 3.76 Å/pix
10,117,438 particles
2x 2D classification, 200 class, uncertainty factor = 4, 40 iterations, batch size 200
8,369,541 re-sorted
1,747,897 selected
605,078 selected
7,764,463 excluded
Ab-initio reconstruction
Ab-initio reconstruction
1,491,319 particles
Extract from micrographs: Box = 144 pix, 1.88 Å/pix
Ab-initio reconstruction: 3 classes
1,084,712 particles
Ab-initio reconstruction: 2 classes, $Res_{initial}$ = 9 Å, $Res_{max}$ = 7 Å
799,516 particles
Extract: Box = 288 pix, 0.94 Å/pix
Non-uniform refinement
3.0 Å
Mask generation
P2RY8
7TM
Local refinement:
Gaussian priors
2° SD
1 Å SD
Re-center rot.
Re-center shift
2.9 Å
3D Classification: 4 classes, res = 3.00 Å, hard classification
26%
27%
25%
22%
Remove duplicates: $d_{min}$ = 1 Å
Local refinement: Gauss. priors, 1° and 1 Å SD, Re-center rot. and shift, 7TM mask
3D Classification: 4 classes, res = 2.75 Å, hard classification
28%
1%
29%
43%
90,243 particles
Non-uniform refinement:
(d)
2.9 Å
Local refinement:
Gaussian priors
1° SD
1 Å SD
Re-center rot.
Re-center shift
7TM mask
(e)
2.7 Å
d
Fourier Shell Correlation
Spatial frequency ($Å^{-1}$)
None (3.4 Å)
Loose (3.1 Å)
Tight (2.9 Å)
Corrected (2.9 Å)
Number of images
Elevation (rad)
Azimuth (rad)
e
Fourier Shell Correlation
Spatial frequency ($Å^{-1}$)
None (3.9 Å)
Loose (3.0 Å)
Tight (2.7 Å)
Corrected (2.7 Å)
Number of images
Elevation (rad)
Azimuth (rad)

**Extended Data Fig. 7: Cryo-EM data processing for P2RY8-G13. a,** Representative cryo-electron micrograph from the curated P2RY8-$G_{13}$ data set (n = 12,053 obtained from a Titan Krios microscope). **b,** A subset of highly populated, reference-free 2D-class averages are shown. Scale bar is 50 Å. **c,** Schematic showing the image processing workflow for P2RY8-$G_{13}$. Initial processing was performed using UCSF MotionCor2 and cryoSPARC, where particles were sorted using a combination of 2D classification, ab-inition reconstruction, and 3D classification. Finally EM maps were obtained in cryoSPARC using the non-uniform and local refinement tools. Dashed boxes indicated selected classes, and 3D volumes of classes and refinements are shown along with the global gold-standard Fourier shell correlation (GSFSC) resolutions. **d, e,** Map validation for the P2RY8- $G_{13}$, **d,** globally refined, and **e,** locally refined cryo-EM maps. GSFSC curves are calculated in cryoSPARC. Euler angle distributions calculated in cryoSPARC are also provided for each map.

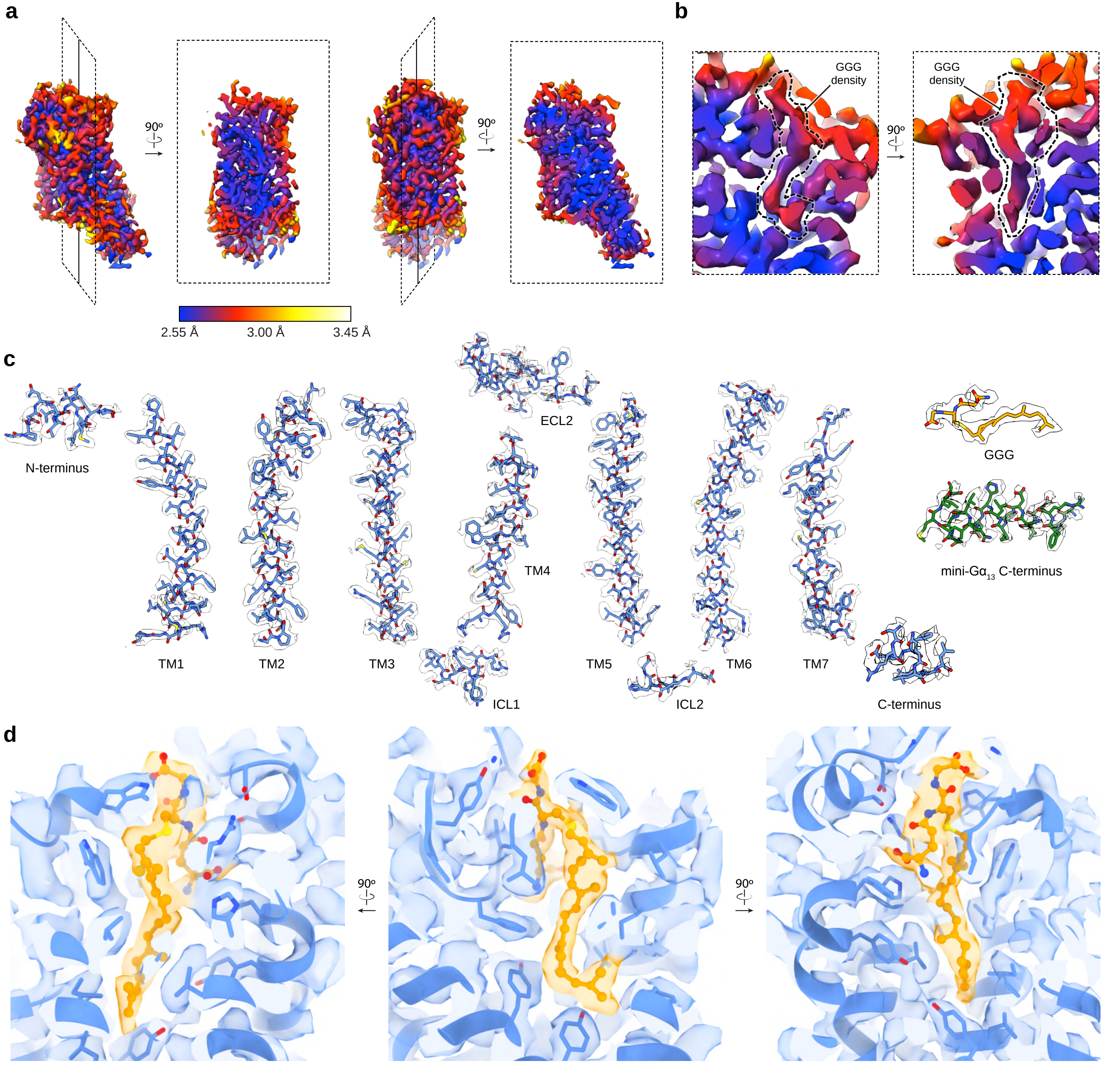

a
90º
90º
2.55 Å
3.00 Å
3.45 Å
b
GGG density
90º
GGG density
c
N-terminus
TM1
TM2
TM3
ECL2
TM4
ICL1
TM5
ICL2
TM6
TM7
C-terminus
GGG
mini-Gα13 C-terminus
d
90º
90º

**Extended Data Fig. 8: Cryo-EM density and atomic model. a,** Orthogonal views of local resolution for the locally refined map covering the 7TM domain of the P2RY8-$G_{13}$ complex, calculated with the local resolution estimation tool in cryoSPARC. **b,** Close-up view showing the local resolution of the GGG binding site. **c,** Representative cryo-EM densities from the 3D reconstruction of P2RY8 from a sharpened, locally refined map of P2RY8-$G_{13}$ at a map threshold of 1.02. Shown are the transmembrane helices and loop regions of P2RY8, the C-terminal helix of mini-$G\alpha_{13}$, as well as GGG. **d,** Close-up views of cryo-EM density supporting GGG binding pose (orange sticks and density) using a sharpened, locally refined map of P2RY8-$G_{13}$ at a map threshold of 1.02.

#### Extended Data Figure 9

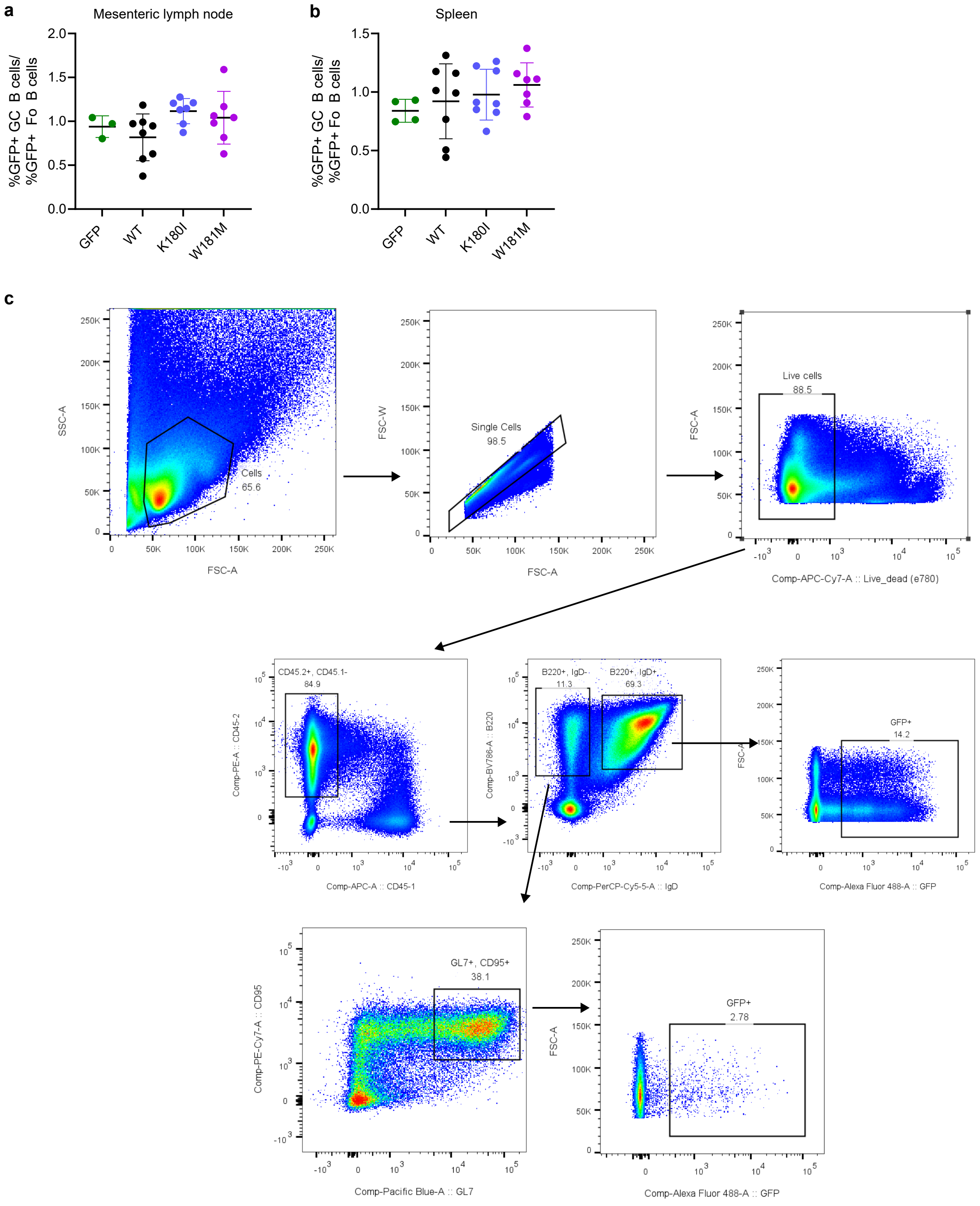

**Extended Data Fig. 9: In vivo variant analysis in spleen and mesenteric lymph node. a,b** Irradiated CD45.1 mice were reconstituted with bone marrow transduced with EV-GFP, WT-P2RY8-GFP, K180I-GFP, or W181M-GFP. After reconstitution, mesenteric lymph nodes (**a**) and spleen (**b**) were analyzed for the frequency of GFP+ cells among GC and follicular B cells and the ratio plotted. **c,** Representative flow cytometry gating; this depicts a Peyer's patch sample. GC B cells identified as singlet live cells, $B220^+$, $IgD^-$, $GL7^+$, $CD95^+$; follicular B cells identified as singlet live cells, $B220^+$, $IgD^+$. **a,b** Pooled from 3 experiments, each point is a mouse. Graphs show means and SDs.

#### Extended Data Figure 10

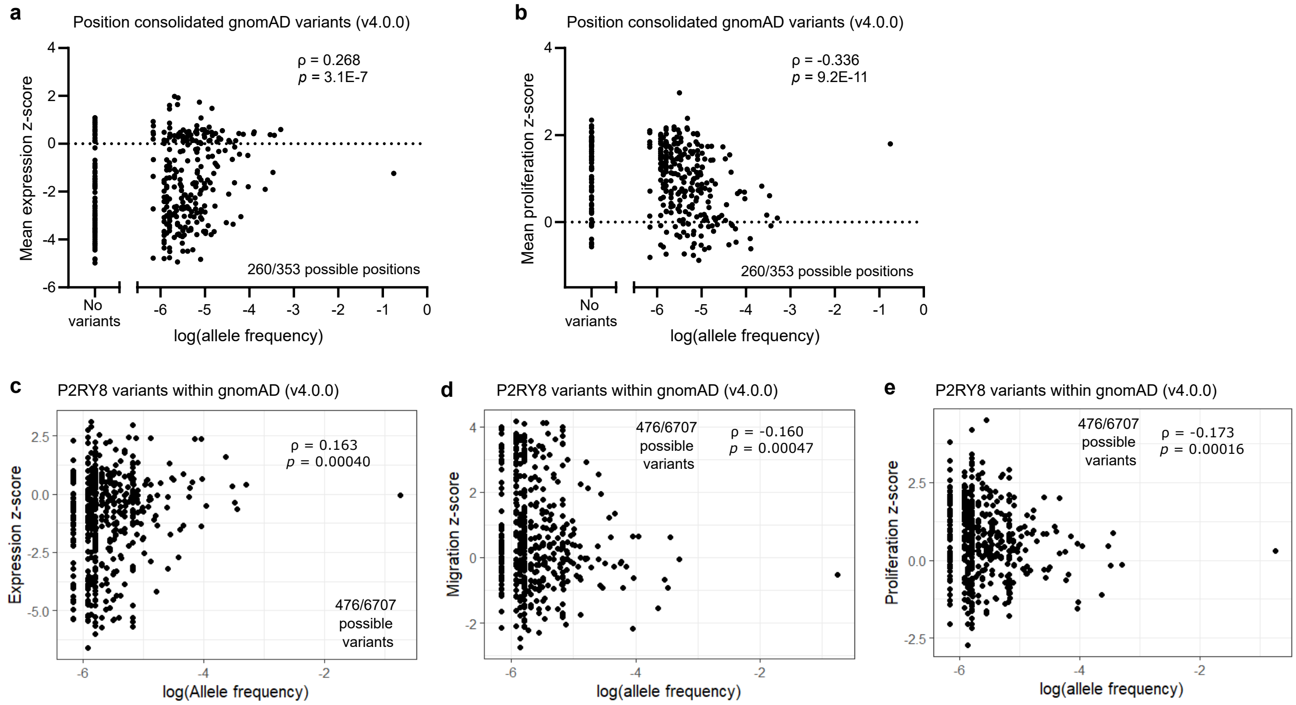


**f**

Expression

| | gnomAD | DLBCL/BL | Non-hem COSMIC |
|---|---|---|---|
| All var | D=0.2039 p=2.22E-16 | D=0.3223 p=1.51E-4 | D=0.2222 p=5.81E-7 |
| Non-hem COSMIC | D=0.153 p=0.0081 | D=0.534 p=6.69E-10 | |
| DLBCL/BL | D=0.5205 p=2.76E-10 | | |

Migration

| | gnomAD | DLBCL/BL | Non-hem COSMIC |
|---|---|---|---|
| All var | D=0.2027 p=4.44E-16 | D=0.3795 p=3.87E-6 | D=0.2316 p=1.59E-7 |
| Non-hem COSMIC | D=0.1963 p=2.39E-4 | D=0.5839 p=7.31E-12 | |
| DLBCL/BL | D=0.5724 p=2.37E-12 | | |

Proliferation

| | gnomAD | DLBCL/BL | Non-hem COSMIC |
|---|---|---|---|
| All var | D=0.2141 p=2.2E-16 | D=0.2971 p=6.29E-4 | D=0.1688 p=3.36E-4 |
| Non-hem COSMIC | D=0.0642 p=0.719 | D=0.4596 p=2.42E-7 | |
| DLBCL/BL | D=0.5023 p=1.31E-9 | | |

**Extended Data Fig. 10: Additional germline and lymphoma-associated P2RY8 variant analysis. a,b,** Plot comparing summed allele frequency for all missense variants within each position in gnomAD (v4.0.0) to **(a)** mean expression z-score or **(b)** mean proliferation z-score for each position. **c-e,** Plot comparing missense variant allele frequency in gnomAD (v4.0.0) to the **(c)** expression, **(d)** migration, or **(e)** proliferation z-scores for those variants. **f,** Tables of Kolmogorov-Smirnov test results comparing expression, migration, and proliferation score distributions for all missense variants, variants present in gnomAD (v4.0.0), non-hematologic cancer variants from COSMIC, and set of DLBCL and Burkitt lymphoma variants. Asymptotic two-sample Kolmogorov-Smirnov test performed; p-values not adjusted for multiple comparison. **a-e,** ρ, Spearman correlation; p-value calculated using algorithm AS 89 with Edgeworth series approximation.

**Supplemental Tables:**

**Supplemental Table 1: A**ll variants in library with frequency data supplied by Twist; also gnomAD variants, lymphoma variants, and non-hematologic cancer P2RY8 variants analyzed in this study.

**Supplemental Table 2:** DMS results; sheet 1 contains GATK output variant counts; sheets 2-4 contain Enrich2 scores; sheet 5 contains normalized variant data; sheet 6 contains data summarized by position.

**Supplemental Table 3:** Cryo-EM statistics

**Supplemental Table 4:** Primers used in cloning, sequencing library generation, and site-directed mutagenesis.
