## Supplemental Table 3 for "Phenotypic pleiotropy of missense variants in human B cell-confinement receptor P2RY8"

**Table S3. Cryo-EM data collection, refinement, and validation statistics**

| EMDB: Full map  EMDB: 7TM map  RCSB PDB: Model | **GGG-bound**  **P2RY8-G_13_**  EMD-47912  EMD-47914  9ECJ |
| --- | --- |

| **Data collection** |  |
| --- | --- |
| Microscope | Thermo Scientific Krios G3i |
| Detector | Thermo Scientific Falcon 4i with Selectris X energy filter |
| Voltage (kV) | 300 |
| Magnification | 130,000 |
| Defocus range (µm) | -0.8 to -2.1 |
| Pixel size, physical (Å) | 0.94 |
| Total exposure (e^-^/Å^2^) | 50 |
| Images, number of | 12,053 |
| EER fractions | 80 |
| Initial particles, number of | 10,117,438 |
| Final particles, number of | 90,243 |
| Symmetry imposed | C1 |
| Map sharpening, *B* factor (Å^2^)  Full map  7TM map | -79.5  -83.0 |
| Map resolution, masked (Å)  Full map  7TM map | 2.9  2.7 |
| FSC threshold | 0.143 |
| **Refinement** |  |
| Initial model used (AlphaFold code) | Q86VZ1 |
| Model resolution (Å) | 4.6 |
| FSC threshold | 0.5 |
| Model composition |  |
| Chains | 4 |
| Non-hydrogen atoms | 6,330 |
| Protein residues | 790 |
| Ligands | 1 |
| *B* factors (Å^2^) |  |
| Protein | 65.97 |
| Ligand | 20.00 |
| R.m.s. deviations |  |
| Bond length (Å) | 0.017 |
| Bond angles (°) | 2.104 |
| Validation |  |
| MolProbity score | 2.71 |
| Clash score | 11.88 |
| Rotamer outliers (%) | 8.09 |
| Ramachandran plot |  |
| Favored (%) | 93.33 |
| Allowed (%) | 6.67 |
| Disallowed (%) | 0.00 |
